# Dissociable neural information dynamics of perceptual integration and differentiation during bistable perception

**DOI:** 10.1101/133801

**Authors:** Andrés Canales-Johnson, Alexander J. Billig, Francisco Olivares, Andrés Gonzalez, María del Carmen Garcia, Walter Silva, Esteban Vaucheret, Carlos Ciraolo, Ezequiel Mikulan, Agustín Ibanez, David Huepe, Srivas Chennu, Tristan A. Bekinschtein

**Author notes:** Correspondence: Andres Canales-Johnson, PhD, Department of Psychology, University of Cambridge, Downing Site, CB2 3EB, Cambridge, United Kingdom. **Author’s contributions** Conceived and designed the experiments: ACJ, TAB. Performed the experiments: ACJ, FO, AG, EM. Analysed the data: ACJ. Contributed reagents/materials/analysis tools: AB, DH, SC. Gave access to clinical patient: MCG, WS, EV, CC. Wrote the paper: ACJ, AB, AI, SC, TAB.

## Abstract

At any given moment, we experience a perceptual scene as a single whole and yet we may distinguish a variety of objects within it. This phenomenon instantiates two properties of conscious perception: integration and differentiation. Integration to experience a collection of objects as a unitary percept, and differentiation to experience these objects as distinct from each other. Here we evaluated the neural information dynamics underlying integration and differentiation of perceptual contents during bistable perception. Participants listened to a sequence of tones (auditory bistable stimuli) experienced either as a single stream (perceptual integration) or as two parallel streams (perceptual differentiation) of sounds. We computed neurophysiological indices of information integration and information differentiation with electroencephalographic and intracranial recordings. When perceptual alternations were endogenously driven, the integrated percept was associated with an increase in neural information-integration and a decrease in neural differentiation across frontoparietal regions, whereas the opposite pattern was observed for the differentiated percept. However, when perception was exogenously driven by a change in the sound stream (no bistability) neural oscillatory power distinguished between percepts but information measures did not. We demonstrate that perceptual integration and differentiation can be mapped to theoretically-motivated neural information signatures, suggesting a direct relationship between phenomenology and neurophysiology.

## INTRODUCTION

Phenomenologically, conscious experience does not only depend on the external information we receive from the environment, but also on internal information that is independent of sensory stimulation. This dissociation between external stimulation and conscious experience is observed in several visual and auditory perceptual illusions in which two or more internally driven percepts alternate under unchanging external stimulation (Sterzer et al. 2009). Moreover, conscious experience cannot be divided into discrete independent components (i.e., it is perceptually integrated), but it can contain an assortment of events and objects (i.e., it is perceptually differentiated) (Tononi et al. 2016). Thus, while integration is the property of experiencing a collection of objects as a unitary percept, differentiation is the property of experiencing these objects as distinct from each other. What are the neural markers of the integration and differentiation of internally driven perceptual contents? We propose that integration and differentiation of internally driven percepts can be neurophysiologically investigated during auditory bistability. During the particular form of auditory bistability employed here, an invariant sequence of tones is experienced as forming either an integrated percept (one stream) or a differentiated percept (two streams) (Snyder et al. 2012) (Figure 1).

**Figure 1.**
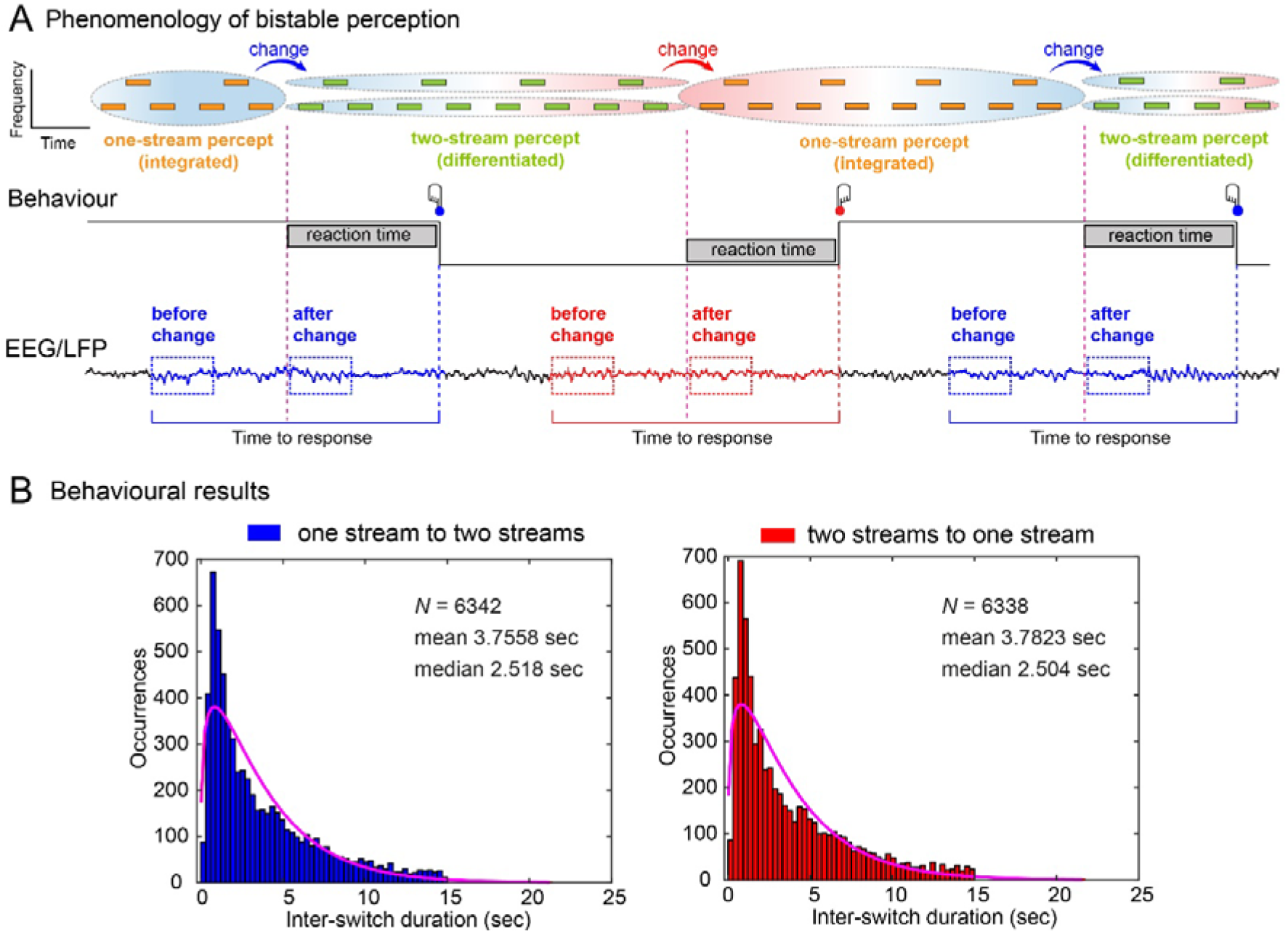
Experimental design and analysis of ongoing brain activity. **(A)** *Top row*: Phenomenology (reported experience) during auditory bistability. Participants listened to a series of tones of two different frequencies separated by a temporal gap (see Experimental Procedures). Tones are experienced either as one stream (phenomenologically integrated percept; orange blocks surrounded by one ellipse) or as two streams (phenomenologically differentiated percept; green blocks surrounded by two ellipses). Perceptual transitions occur either in the one-stream to two-stream direction (blue arrows and blue background) or in the two-stream to one-stream direction (red arrow and red background). *Middle row*: behavioural responses during the task. Participants pressed one button when perceiving that one stream had fully changed into the two-stream percept (blue button) and another button when perceiving that two streams had fully changed into the one-stream percept (red button). *Bottom row*: dynamical analyses and windows of interest for EEG and iEEG signal analyses. Ongoing activity in during both transitions was calculated using a sliding window procedure over a fixed time window locked to the onset of the button press. Window size was calculated based on the mean reaction times (RT) in the exogenous condition (1342 ms), and the minimum duration between responses that guaranteed no overlap between epochs (2500 ms) (see Materials and Methods). (B) *Left panel*: Inter-switch duration histograms for the one- to two-stream perceptual stable period and for the two- to one-stream perceptual stable period (*Right panel*).

Neurophysiologically, conscious experience is thought to require the joint presence of information integration and information differentiation (Fahrenfort et al., 2012; Oizumi et al. 2014; Tononi et al. 2016; Fahrenfort et al., 2017). In particular, the emergence of conscious percepts is believed to involve the integration of information coming from frontal and parietal brain areas to form a phenomenologically unified whole (Dehaene and Changeux 2011; Dehaene et al. 2014; Tononi et al. 2016). Therefore, a reasonable assumption is that the integrated percept (one stream, in the case of this experiment) should be associated with correspondingly higher neural information integration (NII). Recently, NII has been empirically measured in a direct manner by computing the amount of information shared between long-distance EEG signals, and it has been used to discriminate between vegetative and minimally conscious patients (King et al. 2013; Sitt et al. 2014). This NII measure can detect non-oscillatory coupling between signals as compared to classical measures of neural oscillatory integration (NOI) such as phase synchronization. In the case of auditory bistability, we expect higher neural information integration for the perceptually integrated (one-stream) percept compared to the perceptually differentiated (two-stream) percept, as the former would require information about tones of two different frequencies to form a single, integrated percept.

Complementary to NII, empirical indices of neural information differentiation (NID) have been used to separate levels of consciousness by estimating the degree of compressibility of EEG signals (Casali et al. 2013; Sitt et al. 2014; Schartner et al. 2015, 2017). For instance, a decrease in NID has been observed in patients in vegetative states compared to minimally conscious states (Sitt et al. 2014), showing that differentiation of neural information is associated with a cognitively more advanced state of consciousness, as clinically defined. On the other hand, the only study providing a preliminary indication that neurophysiological differentiation might be related to perceptual processes is an fMRI study (Boly et al. 2015), showing that NID was highest when participants watched a coherent movie, intermediate when scenes were scrambled, and minimal for ‘TV noise’. However, it is unclear whether neurophysiological differentiation was specifically related to conscious awareness since factors such as low-level visual processing, expectations and top-down attention might have influenced the differences observed between conditions. During auditory bistability, we can specifically evaluate perceptual differentiation directly since what is changing is not the stimulus itself but how it is subjectively experienced. If the neural information associated with a conscious percept is highly differentiated, NID is expected to be high since information should be less compressible. In contrast, neural differentiation is expected to be low if EEG signals are processing information in a stereotypical way because information is highly redundant (easily compressed). Following this rationale, we expect that the differentiated (two-stream) percept should be associated with higher neural information differentiation.

In addition, there is ample and rapidly growing evidence that endogenous or ‘ongoing’ brain activity in the gamma band (30-70 Hz) is neither meaningless nor random but instead carries functional information largely determining the way incoming stimuli are interpreted (Engel et al. 2001, 2013; Varela et al. 2001; Freeman 2015). For instance, studies of the visual system have shown that neural oscillatory integration (NOI) in the gamma band is involved in the alternation between visual conscious percepts (Doesburg et al. 2005; Hipp et al. 2012). Thus, drawing upon these results and a wealth of previous research that has identified gamma band activity as relevant for conscious perception (Melloni et al. 2007; Engel et al. 2013; Levy et al. 2015), here we analyse information and oscillatory dynamics of ongoing activity in the gamma band. However, we specifically evaluate the theoretical prediction (Koch et al. 2016) that information dynamics (NII and NID) but not oscillatory dynamics (NOI) of ongoing activity underpin phenomenological integration and differentiation.

By measuring high-density scalp EEG and intracranial EEG (iEEG) in humans, we tested the hypothesis that during the formation of internally driven percepts, conscious experience goes along with neurophysiological indices of information processing. Specifically, we predicted that the perceptually integrated content would correspond to high frontoparietal neural information integration in the gamma band and conversely, the perceptually differentiated content would be associated with high neural information differentiation within frontal and parietal regions.

## Materials and Methods

### Healthy participants and patient

Twenty-nine right-handed healthy participants (14 males; mean ± SD age = 21.30 ± 2.2 years) and one left-handed epileptic patient (female; 29 years) gave written informed consent to take part in the experiment. The study was approved by the institutional ethics committee of the Faculty of Psychology of Universidad Diego Portales (Chile) and the Institutional Ethics Committee of the Hospital Italiano de Buenos Aires, Argentina, in accordance with the Declaration of Helsinki. The patient suffered from drug-resistant epilepsy from the age of 8 years and was offered surgical intervention to alleviate her intractable condition. Drug treatment at the time of implantation included 600 mg/d oxcarbazepine, 200 mg/d topiramate, and 750 mg/d levetiracetam. Computed tomography (CT) and magnetic resonance imaging (MRI) scans were acquired after insertion of depth electrodes. The patient took part in the current study one day before the surgery. She was attentive and cooperative during testing, and her cognitive performance before and one week after the implantation was indistinguishable from healthy volunteers. The patient was specifically recruited for this study because she was implanted with electrodes covering frontal and parietal regions (Table 1). Healthy participants and the epileptic patient performed the task with eyes closed in a dimly lit room.

**Table 1.** Coordinates and anatomical loci of intracranial electrodes analysed.

| Electrode | MNI Coordinates | Cortical region (gyrus) |
| --- | --- | --- |
| 1 | -26; -44; 62 | Left Superior Parietal Lobe |
| 2 | -28; -44; 62 | Left Superior Parietal Lobe |
| 3 | -30; -44; 62 | White Matter |
| 4 | -14; 32; 40 | Left Middle Frontal Gyrus |
| 5 | -17; 32; 40 | Left Middle Frontal Gyrus |
| 6 | -20; 32; 40 | Left Middle Frontal Gyrus |
| 7 | -12; 32; 40 | White Matter |
| 8 | -8; 32; -10 | Left Orbitofrontal Cortex |
| 9 | -11; 32; -10 | Left Orbitofrontal Cortex |
| 10 | -14; 32; -10 | Left Orbitofrontal Cortex |
| 11 | -16; 32; 10 | White Matter |
| 12 | -26; -28; 62 | Left Primary Motor Cortex |
| 13 | -13; -28; 62 | White Matter |
| 14 | -40; -8; 4 | Left Posterior Insular Cortex |
| 15 | -40; -30; 4 | White Matter |

Note: Electrodes are depicted in Figure 4 and Supplementary Figures 8 to 13. SPL = Superior Parietal Lobe. MFG = Middle Frontal Gyrus. OF = Orbitofrontal Cortex. MC = Motor Cortex. PIC = Posterior Insular Cortex.

### Stimuli and experimental conditions

There were two experimental conditions. In the endogenous condition (bistability), we used a bistable auditory tone sequence (Carlyon et al. 2001; Gutschalk et al. 2005; Pressnitzer and Hupé 2006). Participants listened to a repeating pattern of three tones (ABA), each group of which was followed by a silence (‘-’); the pattern is experienced either as a one-stream percept or as a two-stream percept (Figure 1A). The frequency of the A tone was 587 Hz and that of the B tone was 440 Hz (5 semitones difference). The duration of each tone was 120 ms. The silence (‘-’) that completed the ABA… pattern was also 120 ms long, thus making both the set of A tones and the set of B tones isochronous (Pressnitzer and Hupé 2006). Participants were instructed to press a button with one hand when perceiving that the one-stream percept had fully changed into two streams and a second button with the other hand when perceiving that the two-stream percept had fully changed into one stream (Figure 1A, middle panel).

In the exogenous condition (control), participants listened to repeating patterns of three (ABA) or two (AB) tones, each group of which was separated by one (‘-’) or two periods of silence (‘--’) respectively. In both cases the A and B tones had the same parameters (frequencies and duration) and the silence had the same duration as in the endogenous condition (ABA- pattern). In the exogenous condition blocks of each pattern (ABA- and AB--) were constructed to last 4-8 seconds before changing to the other pattern. This suppressed the effect of endogenous bistability, such that ABA- was most often perceived as one stream, whereas AB-- was largely perceived as two streams. As in the endogenous conditions, participants were instructed to press a button with one hand when perceiving that ABA- had fully changed into AB-- and another button with the other hand when pattern AB-- had fully changed into ABA-. Importantly, the exogenous condition allowed us to characterize the dynamics of neural activity specifically related to external changes in the stimuli (the two alternating patterns) and to contrast them with the dynamics of internal neural activity elicited by the endogenous condition (bistability). The endogenous and exogenous conditions used physically similar stimuli, with the latter sometimes having one fewer A tone. Because the analysis windows were not time-locked to the stimuli, but rather to responses (see “Analysis of ongoing neural activity”), differences in the evoked responses to specific tones are unlikely to account for the observed pattern of results.

In order to match both conditions as closely as possible, the exogenous condition was always preceded by the endogenous condition. By designing the experiment in this manner, we were able to quickly calculate the number of both types of alternations during the endogenous conditions on a participant-per-participant basis. Based on this individual calculation, we structured the subsequent exogenous condition based on the number of switches of the endogenous condition. Similarly, the pattern of alternation (i.e. stimuli sequence) in the exogenous conditions was defined based on the first switch observed during the endogenous condition (i.e. the sequence started either with a one-stream stimulus or with a two-stream stimulus). This procedure was meant to control for differences in the number of alternations and for differences in the sequence of alternations between conditions.

### Electroencephalography (EEG) recordings, pre-processing and analysis

EEG signals were recorded with 128-channel HydroCel Sensors using a GES300 Electrical Geodesic amplifier at a sampling rate of 500 Hz using the NetStation software. During recording and analyses, the electrodes’ average was used as the reference electrode. Two bipolar derivations were designed to monitor vertical and horizontal ocular movements. Following Chennu *et al* (Chennu et al. 2014), data from 92 channels over the scalp surface were retained for further analysis. Channels on the neck, cheeks and forehead, which reflected more movement-related noise than signal, were excluded. Eye movement contamination, cardiac and muscle artefacts were removed from data before further processing using an independent component analysis (ICA) (Delorme and Makeig 2004). All conditions yielded at least 91% of artefact-free trials. Trials (−2500 to 0 ms relative to a button press) that contained voltage fluctuations exceeding ± 200 μV or transients exceeding ± 100 μV were also excluded. The EEGLAB MATLAB toolbox was used for data pre-processing and pruning (Delorme and Makeig 2004). MATLAB open source software FieldTrip (Oostenveld et al. 2011) and customized scripts were used for calculating Neural Information Integration (NII), Neural Information Differentiation (NID), Neural Oscillatory Integration (NOI) and Neural Oscillatory Power (NOP) measures. No filtering was performed during the pre-processing stage.

### Intracranial electroencephalography (iEEG): recordings and pre-processing

Stereo-electroencephalography (SEEG), an intracranial electroencephalography (iEEG) recording technique in which depth electrodes are inserted in the brain of epileptic patients, were obtained from semi-rigid, multi-lead electrodes implanted in a single patient. The electrodes had a diameter of 0.8 mm and consisted of 5, 10 or 15 contact leads that were 2-mm wide and 1.5-mm apart (DIXI Medical Instruments). The electrode shafts were implanted in different regions of the frontal, temporal, central and parietal cortices and subcortical structures. For the purposes of the current study, iEEG recordings were analysed from the left orbitofrontal cortex (OF), left middle frontal gyrus (MFG), left superior parietal lobe (SPL), left primary motor cortex (MC) and left posterior insular cortex (PIC) (Table 1). Each electrode located in grey matter was re-referenced to an electrode of the same electrode shaft located in white matter (Li et al., 2018). Following this procedure, we preserved the limited number of electrodes in SPL (2 electrodes), OF (3 electrodes) and MFG (3 electrodes) that would have been reduced in number using a bipolar reference (N−1). MNI coordinates of the depth electrodes were obtained from MRI and CT images using SPM (Friston 2006) and MRIcron (Rorden and Brett 2000) software. The recordings were sampled at 1024 Hz and down-sampled to 500 Hz for further analysis. The exact MNI coordinates and cortical regions of the selected electrodes are reported in Table 1. Open-source BrainNet Viewer software was used for visualization of selected electrodes (Xia et al. 2013).

### Analysis of ongoing neural activity

A classical experimental approach for studying endogenous, or “ongoing” activity in the EEG related to internal fluctuations during cognitive tasks is by analysing the EEG window before the onset of motor responses when participants report internal changes. This approach has been used for studying neural signatures of conscious awareness, such as in bistable perception (Parkkonen et al. 2008), binocular rivalry (Doesburg et al. 2005; Frässle et al. 2014) and intrusions of consciousness (Noreika et al. 2015). Here, ongoing EEG and iEEG activity (not time-locked to stimuli) preceding the onset of each response (button press), and presumed to span the change in perception, was analysed in terms of connectivity (NII and NOI) and complexity (NII).

Window size selection was based on the following procedure. First, we calculated the mean reaction time in the exogenous condition (M = 1342 ms; SD = 101 ms). Second, in the endogenous condition, we calculated the minimum inter-switch duration such that windows would not overlap after EEG data epoching (the minimum window size was 2500 ms; epoching from −2500 to 0 ms relative to button press). Third, as the mean RT in the exogenous condition (1342 ms) was approximately half of the window size of the endogenous condition (2500 ms), we selected a window of analysis of 2500 ms for both conditions (from −2500 to 0 ms relative to button press) (Figure 1, lower panel). Importantly, this window included the onset of the exogenous auditory patterns (‘ABA-’ and ‘AB--’). The same procedure was repeated for the intracranial patient (reaction times in the exogenous condition: M = 1380 ms, SD = 79 ms; window size in the endogenous conditions: 2500 ms).

For statistical analyses (see below), two time windows of 500 ms were selected based on the exogenous condition. A window after the change between auditory percepts (after-change window (AC)) was defined based on the mean reaction time at the group level (from −1342 to −842 ms). A second time window was defined at the epoch onset (before-change window (BC); from −2500 to −2000 ms). The rationale behind the latencies of both time windows was to select *a priori* the periods when both externally driven percepts remained perceptually stable. Windows were not selected using a data-driven approach (see Supplementary Figure 1). The same window lengths and latencies were used for the endogenous condition (Figure 1).

### Neural oscillatory power (NOP): spectral power

In order to derive our measure of neural oscillatory power, we first band-pass filtered the raw signal using a 6th order Butterworth filter. Spectral power was estimated by calculating the square of the envelope obtained from the absolute value of the Hilbert transform after filtering. This procedure provides the instantaneous power emission which reflects the strength of local synchronization. The frequency bands were defined as follows: theta (4–7 Hz), alpha (8–12 Hz), beta (15–25 Hz), gamma (30–60 Hz), following the canonical frequency-band classification (e.g. Li et al., 2018).

### Phase synchronization

We quantified phase coherence between pairs of electrodes as a measure of dynamical linear coupling among signals oscillating in the same frequency band. Phase synchronization analysis proceeds in two steps: (i) estimation of the instantaneous phases and (ii) quantification of the phase locking.

### Estimation of the instantaneous phases

To obtain the instantaneous phases *φ* of the neural signals, we used the Hilbert transform approach (Foster et al. 2016). The analytic signal *ξ*(*t*) of the univariate measure *x*(*t*) is a complex function of continuous time *t* defined as:

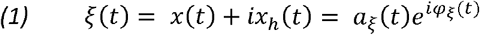

where the function *x*_*h*_(*t*) is the Hilbert transform of *x*(*t*):

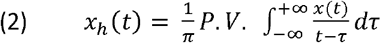

P.V. indicates that the integral is taken in the sense of Cauchy principal value. Sequences of digitized values give a trajectory of the tip of a vector rotating counterclockwise in the complex plane with elapsed time.

The vector norm at each digitizing step *t* is the state variable for instantaneous amplitude *a*_*ξ*_(*t*). This amplitude corresponds to the length of the vector specified by the real and imaginary part of the complex vector computed by Pythagoras’ law and is equivalent to the magnitude of the observed oscillation at a given time and frequency point.

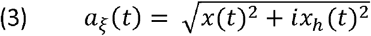

and the arctangent of the angle of the vector with respect to the real axis is the state variable for instantaneous phase *φ*_*x*_(*t*).

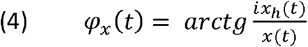

The instantaneous phase *φ*_*x*_(*t*) of *x*(*t*) is taken equal to *φ*_*ξ*_(*t*). Identically, the phase *φ*_*y*_(*t*) is estimated from *y*(*t*). This phase is thus the angle of the vector specified by the real and imaginary components. For a given time and frequency point, it corresponds to a position inside the oscillation cycle (peak, valley, rising, or falling slope).

The instantaneous phase, although defined uniquely for any kind of signal to which the Hilbert transform can be applied, is difficult to interpret physiologically for broadband signals. For this reason, a standard procedure is to consider only narrow-band phase synchronization by estimating an instantaneous phase for successive frequency bands, which are defined by band-pass filtering the time series (Le Van Quyen et al. 2001). Thus, for each trial and electrode, the instantaneous phase of the signal was extracted at each frequency of the interval 1- 60 Hz (in 1-Hz steps) by computing the Hilbert transform using a zero phase shift non-causal finite impulse filter.

### Neural oscillatory integration (NOI): weighted phase lag index (wPLI)

Phase synchronization is often calculated from the phase or the imaginary component of the complex cross-spectrum between the signals measured at a pair of channels. For example, the well-known Phase Locking Value (PLV) (Lachaux et al. 1999) is obtained by averaging the exponential magnitude of the imaginary component of the cross-spectrum. However, such phase coherence indices derived from EEG data are affected by the problem of volume conduction, and as such they can have a single dipolar source, rather than a pair of distinct interacting sources, producing spurious coherence between spatially disparate EEG channels. The Phase Lag Index (PLI), first proposed by Stam *et al* (Stam et al. 2007) attempts to minimize the impact of volume conduction and common sources inherent in EEG data, by averaging the signs of phase differences, thereby ignoring average phase differences of 0 or 180 degrees. This is based on the rationale that such phase differences are likely to be generated by volume conduction of single dipolar sources. However, despite being insensitive to volume conduction, PLI has a strong discontinuity in the measure, which causes it to be maximally sensitive to noise.

The Weighted Phase Lag Index (wPLI) (Vinck et al. 2011) addresses this problem by weighting the signs of the imaginary components by their absolute magnitudes. Further, as the calculation of wPLI also normalises the weighted sum of signs of the imaginary components by the average of their absolute magnitudes, it represents a dimensionless measure of connectivity that is not directly influenced by differences in spectral or cross-spectral power. For these reasons, we employed the wPLI measure to estimate connectivity in our data. The wPLI index ranges from 0 to 1, with value 1 indicating perfect synchronization (phase difference is perfectly constant throughout the trials) and value 0 representing total absence of synchrony (phase differences are random). For each trial and pair of electrodes, wPLI was estimated using a 500 ms sliding window with 2 ms time step, i.e. with a 96% overlap between two adjacent windows.

### Neural information integration (NII): weighted symbolic mutual information (wSMI)

In order to quantify the coupling of information flow between electrodes we computed the weighted symbolic mutual information (wSMI) (King et al. 2013; Sitt et al. 2014). This measure assesses the extent to which two signals present joint non-random fluctuations, suggesting that they share information. wSMI has three main advantages: (i) it allows for a rapid and robust estimation of the signals’ entropies; (ii) it provides an efficient way to detect non-linear coupling; and (iii) it discards spurious correlations between signals arising from common sources, favouring non-trivial pairs of symbols. For each trial, wSMI is calculated between each pair of electrodes after the transformation of the EEG and iEEG signals into a sequence of discrete symbols defined by the ordering of k time samples separated by a temporal separation τ (King et al. 2013). The symbolic transformation depends on a fixed symbol size (k = 3, that is, 3 samples represent a symbol) and a variable τ between samples (temporal distance between samples) which determines the frequency range in which wSMI is estimated (Sitt et al. 2014). In our case, we chose τ = 32 and 6 ms to isolate wSMI in alpha (wSMI_α_) and gamma (wSMI_γ_) bands respectively. The frequency specificity *f* of wSMI is related to k and τ as:

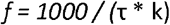

As per the above formula, with a kernel size k of 3, τ values of 32 and 6 ms produced a sensitivity to frequencies below and near to 55 Hz (gamma range) and 10 Hz (alpha range), respectively, with these two frequency values used as low-pass filters in order to avoid aliasing artifacts (King et al., 2013)

wSMI was estimated for each pair of transformed EEG and iEEG signals by calculating the joint probability of each pair of symbols. The joint probability matrix was multiplied by binary weights to reduce spurious correlations between signals, as confirmed by simulations and empirical data in the original wSMI publication (King et al., 2013). The weights were set to zero for pairs of identical symbols, which could be elicited by a unique common source, and for opposite symbols, which could reflect the two sides of a single electric dipole. wSMI is calculated using the following formula:

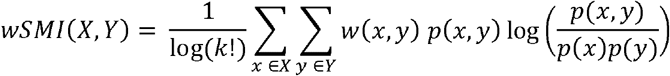

where *x* and *y* are all symbols present in signals *X* and *Y* respectively, w(*x*,*y*) is the weight matrix and *p*(*x*,*y*) is the joint probability of co-occurrence of symbol *x* in signal *X* and symbol *y* in signal *Y*. Finally, *p*(*x*) and *p*(*y*) are the probabilities of those symbols in each signal and *k* is the number of symbols - used to normalize the mutual information (MI) by the signal’s maximal entropy. Temporal evolution of wSMI was calculated using a 500 ms sliding window with 2 ms time step, i.e. with a 96% overlap between two adjacent windows.

In order to further address possible volume conduction issues in our scalp EEG analyses, we compared wSMI with the non-weighted versions of the metric (Symbolic Mutual Information; SMI), computing both metrics as a function of Euclidean inter-electrode distance (Supplementary Figure 7). For distances below 5 cm, wSMI quickly dropped toward zero, as expected given that the weighted version of this measure was designed to eliminate common source artifacts and to provide higher spatial selectivity (King et al., 2013) (Supplementary Figure 7, *upper row*). In contrast, the unweighted version of the metric (SMI) exhibited the highest values below 5 cm, typically produced by common source artifacts (Supplementary Figure 7, *lower row*). These analyses further confirm that the differences observed in our data are most likely due to a non-spurious spatial relationship between electrode pairs.

### Neural information differentiation (NID): Kolmogorov-Chaitin complexity (K complexity)

Kolmogorov-Chaitin complexity quantifies the algorithmic complexity (Kolmogorov 1965; Chaitin 1974) of an EEG signal by measuring its degree of redundancy (Sitt et al. 2014; Schartner et al. 2015, 2017). Algorithmic complexity of a given EEG sequence can be described as the length of shortest computer program that can generate it. A short program corresponds to a less complex sequence. K complexity was estimated by quantifying the compression size of the EEG using the Lempel-Ziv zip algorithm (Lempel and Ziev 1976).

Algorithmic information theory was introduced by Andreï Kolmogorov and Gregory Chaitin as an area of interaction between computer science and information theory. The concept of algorithmic complexity or Kolmogorov-Chaitin complexity (K complexity) is defined as the shortest description of a string (or in our case a time series *X*). That is to say, K complexity is the size of the smallest algorithm (or computer program) that can produce that particular time series. However, it can be demonstrated by *reductio ad absurdum* that there is no possible algorithm that can measure K complexity (Chaitin 1995). To sidestep this issue, we can estimate an upper-bound value of K complexity(*X*). This can be concretely accomplished by applying a lossless compression of the time series and quantifying the compression size. Capitalizing on the vast signal compression literature, we heuristically used a classical open-source compressor gzip (Salomon 2004) to estimate K complexity(*X*). It is important to standardize the method of representation of the signal before compression in order to avoid non-relevant differences in complexity. Specifically, to compute K complexity(*X*):

1. The time series were transformed into sequences of symbols. Each symbol represents, with identical complexity, the amplitude of the corresponding channel for each time point. The number of symbols was set to 32 and each one corresponds to dividing the amplitude range of that given channel into 32 equivalent bins. Similar results have been obtained with binning ranging from 8 to 128 bins (Sitt et al. 2014).
2. The time series were compressed using the compressLib library for Matlab, this library implements the gzip algorithm to compress Matlab variables.
3. K complexity(*X*) was calculated as the size of the compressed variable with time series divided by the size of the original variable before compression. Our premise is that, the larger the size of the compressed string, the more complex the structure of the time series, thus potentially indexing the complexity of the electrical activity recorded at a sensor.

For each trial and channel, K complexity was estimated using a 500 ms sliding window with 2 ms time step, i.e. with a 96% overlap between two adjacent windows.

### EEG electrode cluster analysis and epoch correction

For the hypothesis-driven analyses, canonical bilateral frontal (*n*=26), parietal (*n*=18), and temporal (*n*=12) electrode clusters were selected for spectral power, complexity (K-complexity) and connectivity analysis (wPLI and wSMI) (Supplementary Figure 14). In the case of spectral power and complexity analysis, values within frontal and parietal electrode clusters were averaged per condition and participant. In the case of frontoparietal wPLI, wSMI, we calculated the mean connectivity that every electrode in the frontal cluster shared with every electrode in the parietal cluster. In order to evaluate long-distance interactions between frontal and parietal electrodes, values between pairs of frontal electrodes and between pairs of parietal electrodes were discarded. Similarly, in the case of temporotemporal connectivity analyses, we calculated the mean connectivity that every electrode in the right-temporal cluster shared with every electrode in the left-temporal cluster, discarding connectivity values within pairs of right-temporal electrodes and pairs of left-temporal electrodes. This procedure allowed us to specifically test the role of long-distance interactions (frontoparietal and temporotemporal) during the endogenous and exogenous conditions. Spectral power, K complexity, wSMI and wPLI values of the corresponding regions of interest were averaged per condition and participant. For the exploratory analysis, connectivity between pairs of frontal and pairs of parietal electrodes were analysed separately. In a similar manner as for the hypothesis-driven analysis, connectivity (wSMI) between frontotemporal and temporoparietal electrodes were analysed, within-cluster pairs were excluded.

Since during the endogenous condition there was no true baseline period because we do not know precisely when the spontaneous changes were initiated, after transforming data into complexity (K complexity), connectivity (wPLI and wSMI) and spectral power time series, and creating the corresponding electrode clusters, we subtracted the mean values between −2500 and −700 ms from each data point per epoch and condition. Motor-related activity in the gamma band has been reported ~−200 ms before the button press during auditory bistable perception (Basirat et al. 2008). Thus, although it is common to analyse response-evoked activity by normalizing epochs using the mean of the entire window including the time of the motor response (Doesburg et al. 2005; Fesi and Mendola, 2014; de Jong et al., 2016), we used a more conservative approach by baseline correcting each epoch from −2500 to −700 ms relative to the button press in order to avoid possible contamination due to motor-related artefacts. In order to make both conditions comparable, we performed the same procedure separately on the endogenous and exogenous conditions.

### Statistical analysis

For statistical analysis of wSMI (NII), K complexity (NID), wPLI (NOI) and spectral power (NOD) on the scalp EEG (29 participants) and iEEG data (1 patient), we performed repeated-measures ANOVA (RANOVA) using 3 within-participant factors: condition (endogenous, exogenous), window (before change, after change), and direction (one- to two-stream, two- to one-stream). Bonferroni correction was computed for *post hoc* comparisons, and Bayes Factors of the null and alternative hypothesis are reported (Masson 2011; Jarosz and Wiley 2014). Statistical analyses were performed using Statistical Product and Service Solutions (SPSS, version 20.0, IBM), MATLAB 2018b and open-source statistical software JASP (JASP Team (2017), version 0.8.1.1).

## RESULTS

Twenty-nine healthy participants and one patient implanted with intracranial electrodes listened to a repeating pattern of three tones followed by a temporal gap. This flow of sounds is experienced either as a one-stream percept (perceptual integration) or as a two-stream percept (perceptual differentiation) (Figure 1A, upper panel). Perception tends to alternate between these alternatives every few seconds (Pressnitzer and Hupé 2006). Participants were asked to press a button as they perceived that the one-stream percept had fully changed into the two-stream percept, and a second button when perceiving the two-stream percept had fully changed into the one-stream percept (Figure 1A, middle panel). As an experimental control task, we used a condition where the stimuli were physically manipulated (varying the length of the silence between tones) in order to generate two externally-driven alternating percepts (exogenous condition). Participants had to perform the same task as in endogenous condition. This control allowed us to establish the extent to which neural activity in the endogenous condition was specific to internally driven perceptual switches. It also allowed us to measure typical reaction times to percept changes with known onset times (in the control task), in order to determine suitable time windows for analysis. Furthermore, the exogenous condition allowed us to control for top-down attention in the absence of bistability by creating externally driven switches between percepts as opposed to the internally driven ones (induced endogenously).

### Behavioural results

In the endogenous condition, inter-switch duration (ISD) distributions were approximated well with a gamma distribution (*P*<0.001; Figure 1B), typically reported during bistable perception dynamics (e.g. Parkkonen at al., 2008). Inter-switch duration (ISD) histograms exhibited similar distributions for the one- to two-stream percept and the two- to one-stream percept, with an average dominance duration of 3.75 ± 0.0412 and 3.78 ± 0.0419 seconds respectively (Figure 1B). No significant differences were observed between distributions (*t*_1,28_ = 0.204; *P* = 0.910; Bayes factor (Bf) in favour of the null = 46.19), suggesting that both directions of perceptual change exhibit similar cognitive demand during internally-driven perception.

Similarly, in the case of the exogenous condition, discrimination of the two auditory sequences showed behavioural equivalence as no differences were found in reaction times (RT) between the one- to two-stream percept and the two- to one-stream percept (*t*_1,28_ = 0.204; *P* = 0.840; Bayes factor (Bf) in favour of the null = 4.97).

### Neural information integration (NII) during the endogenous and exogenous conditions

We first investigated the dynamics of neural information integration in the gamma range (NII_γ_). We compared activity during a 500-ms window before a perceptual change with activity after the change (Figure 1A, lower panel; and see Materials and Methods). This approach contrasts with many previous studies of auditory bistability (Gutschalk et al., 2005; Hill et al., 2012; Szalardy et al., 2013; Billig et al. 2018; Sanders et al. 2018), where the analysis windows are time-locked to the stimulus, and allowed us to concentrate on both stable and transition periods of bistable experience. Response-locked analyses focused on how *ongoing* neural activity relates to perception. A repeated-measures ANOVA (RANOVA; see Materials and Methods) revealed a significant triple interaction between condition (endogenous, exogenous), window (before change, after change), and direction (one- to two-stream, two- to one-stream) for NII_γ_ (*F*_1,28_ = 5.73, *P* = 0.024, 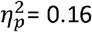, Bayes factor (Bf) in favour of the alternative = 2.73). Bonferroni’s *post hoc* test revealed higher NII_γ_ in the one-stream compared to the two-stream percept in the before-change (BC) window (*P* = 0.009, 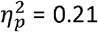, Bf in favour of the alternative = 6.42) (Figure 2A, C). Interestingly, ~1 s later in the after change (AC) window, NII_γ_ again showed higher values for the one-stream than the two-stream percept (*P* = 0.026, 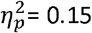, Bf in favour of the alternative = 2.49) (Figure 2A, C). These findings suggest that the phenomenologically integrated percept consistently involved a higher level of gamma neurophysiological integration than the phenomenologically differentiated one.

**Figure 2.**
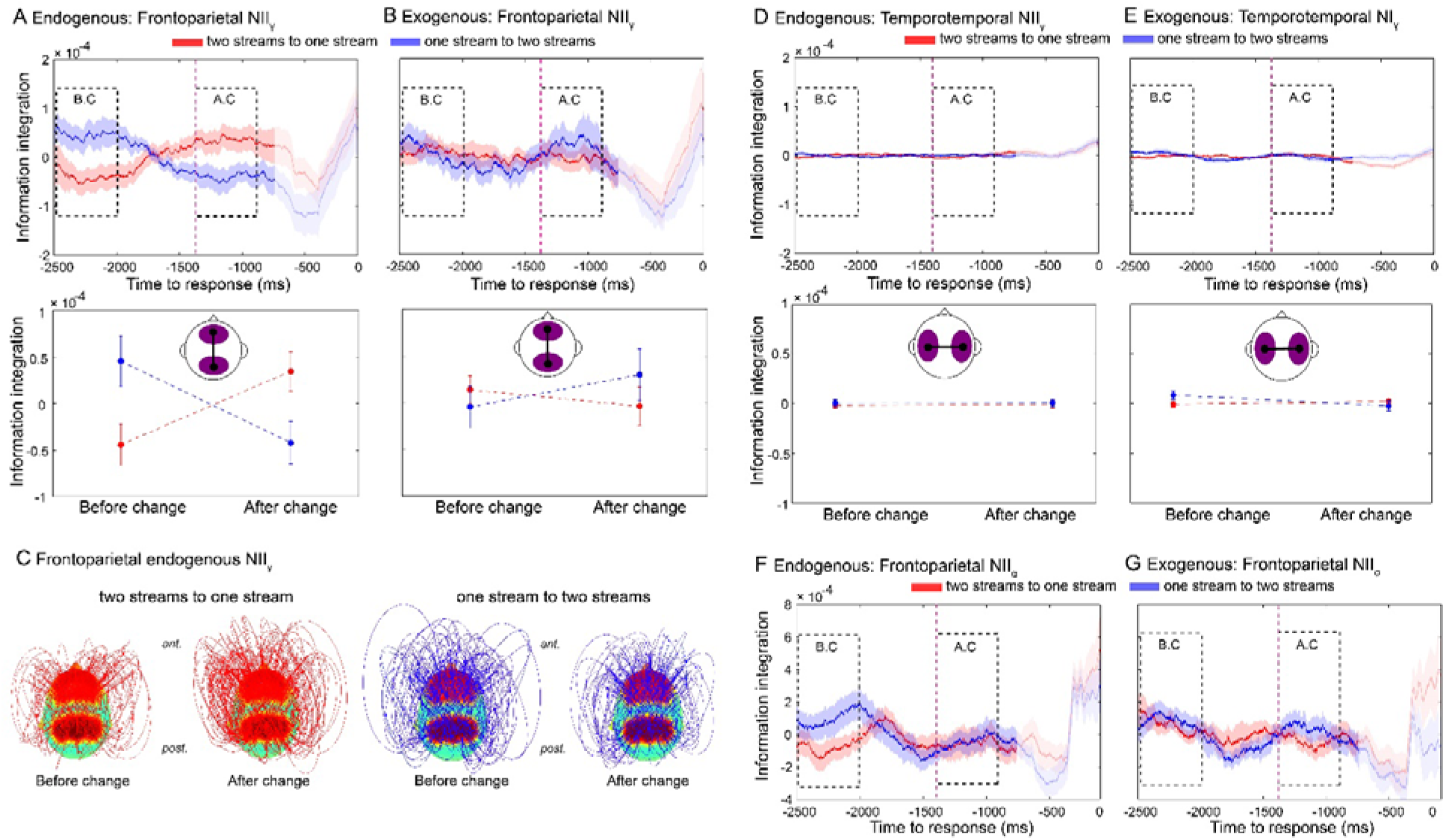
Frontoparietal NII dissociates alternative endogenous percepts during bistable perception. Neural dynamics of NII_γ_ for transitions from the two-stream to the one-stream percept (red line) and from the one-stream to the two-stream percept (blue line) for the endogenous **(A)** and exogenous (control) conditions **(B)**. Purple dashed line marks the mean reaction time (1342 ms) of the exogenous (control) condition. Auditory percepts were directly compared in two windows of interest: the before-change (BC) and the after-change (AC) windows. **(C)** Connectivity topographies for the BC and AC windows for transitions from the two-stream to the one-stream percept (left panel) and from the one-stream to the two-stream percept (right panel) averaged over participants in the endogenous condition. Red areas on the scalp indicate regions of interest (frontal and parietal electrodes, see Experimental Procedures). The height of an arc connecting two nodes indicates the strength of the NII link between them. Values are time-locked to the button press (0 ms) and baseline corrected between −2500 and −700 ms relative to button press (see Materials and Methods). Statistical analyses (bottom row) were computed on two pre-defined 500 ms windows: a BC window (−2500 to 2000 ms) and an AC window (−1342 ms to −842 ms). The onset of both windows was defined based on a control (exogenous) condition in which the stimuli physically change to generate two different percepts (see Experimental Procedures). Shaded bars (top row) and error bars (middle row) represent s.e.m. **(D, E)** Temporotemporal NII in the endogenous and exogenous conditions. **(F, G)** Frontoparietal NII in the alpha band (NII_α_) during endogenous **(F)** and exogenous **(G)** conditions.

Interestingly, while NII_γ_ discriminated between conscious percepts during the endogenous condition (Figure 2A), it did not do so in the exogenous condition (BC: *P* = 0.561; Bf in favour of the null = 5.12 and AC: *P* = 0.349, Bf in favour of the null = 3.26; Bonferroni-corrected *post hoc* tests) (Figure 2B), suggesting that frontoparietal NII_γ_ may be specifically indexing endogenously generated percepts. Furthermore, no differences were observed in the mean level of NII_γ_ between endogenous and exogenous conditions (main effect of condition: *P* = 0.248, Bf in favour of the null = 2.12) or between windows (main effect of window: *P* = 0.918, Bf in favour of the null = 3.59), indicating that the overall amount of information sharing within the frontoparietal network was similar across conditions. This index of information integration was hence sensitive to endogenously driven perceptual changes and may specifically underlie the formation of conscious auditory percepts.

In order to establish the specific role of frontoparietal signals, we computed inter-hemispheric NII_γ_ between temporal electrodes. RANOVA found no reliable triple interaction in temporotemporal NII_γ_ (*F*_1,28_ = 1.51, *P* = 0.228, Bf in favour of the null = 2.02) (Figure 2D, E), implying relatively specific involvement of frontoparietal networks in the emergence of endogenous auditory percepts. Finally, and in addition to NII_γ_, we investigated whether neural information integration in the alpha band (NII_α_) dissociates between auditory percepts, as alpha activity has been previously related to perceptual bistability (Flevaris et al., 2013; Handel and Jensen, 2014). However, the ability of frontoparietal NII to track and distinguish between different endogenous percepts seems to be specifically related to the gamma range since RANOVA revealed no triple interaction between percepts in frontoparietal NII_α_ (*F*_1,28_ = 1.01, *P* = 0.321, Bf in favour of the null = 2.48) (Figure 2F).

Finally, we performed *post hoc* exploratory analysis in order to further investigate NII_γ_ differences between other electrode regions (i.e. frontofrontal, parietoparietal, temporoparietal and frontotemporal pairs of electrodes). RANOVA revealed no triple interaction (condition × window × direction) between these electrode clusters (Supplementary Figure 2). Similarly, we explored differences between these pairs of electrodes in other canonical frequency bands (theta: NII_θ_, alpha: NII_α_, and beta: NII_β_). RANOVA revealed no significant triple interaction (condition × window × direction) for the same groups of electrodes for theta (NII_θ_: Supplementary Figure 3), alpha (NII_α_: Supplementary Figure 4) and beta bands (NII_β_: Supplementary Figure 5). These exploratory results further support the specific role of NII_γ_ in discriminating between internally-driven conscious percepts.

**Figure 3.**
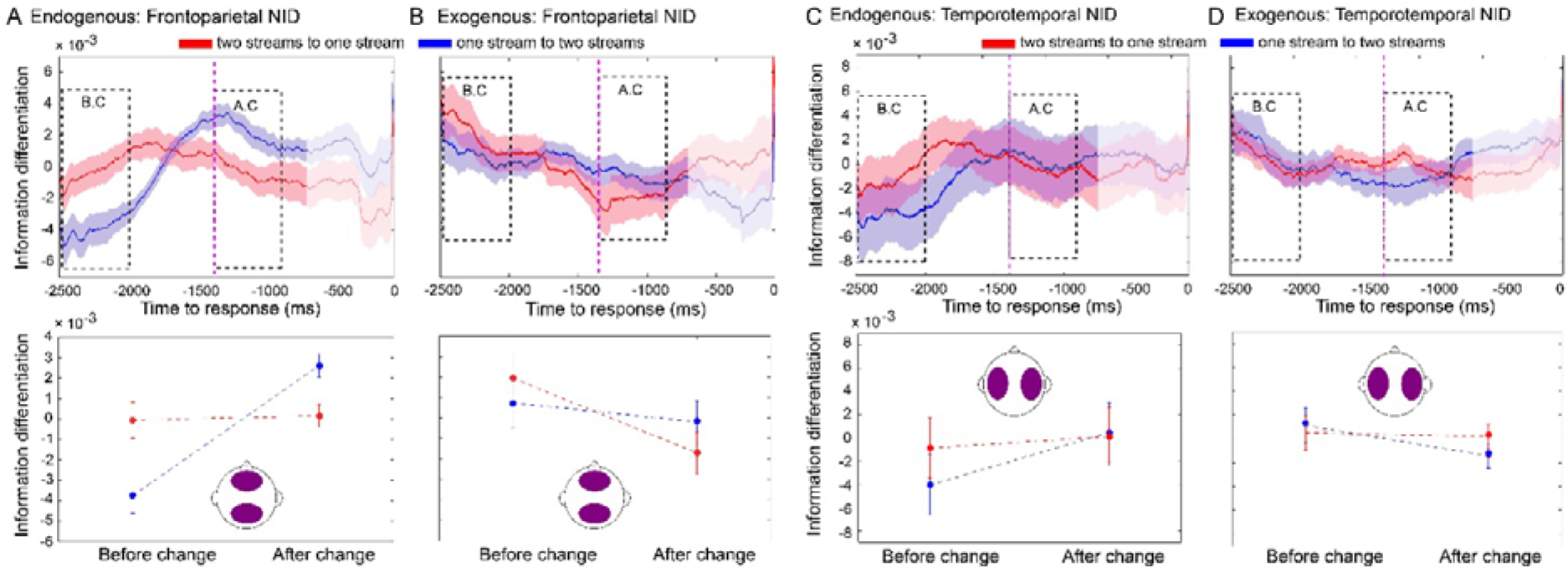
Frontoparietal NID dissociates alternative endogenous percepts during bistable perception. Neural dynamics of NID from the two-stream to the one-stream percept (red line) and from the one-stream to the two-stream percept (blue line) for the endogenous **(A)** and exogenous (control) conditions **(B)**. **(C, D)** Temporotemporal NID dynamics for the endogenous **(C)** and exogenous (control) conditions **(D)**. Purple dashed line marks the mean reaction time (1342 ms) of the exogenous (control) condition. Statistical analysis was performed as described in Figure 2. Shaded bars (top row) and error bars (bottom row) represent s.e.m.

**Figure 4.**
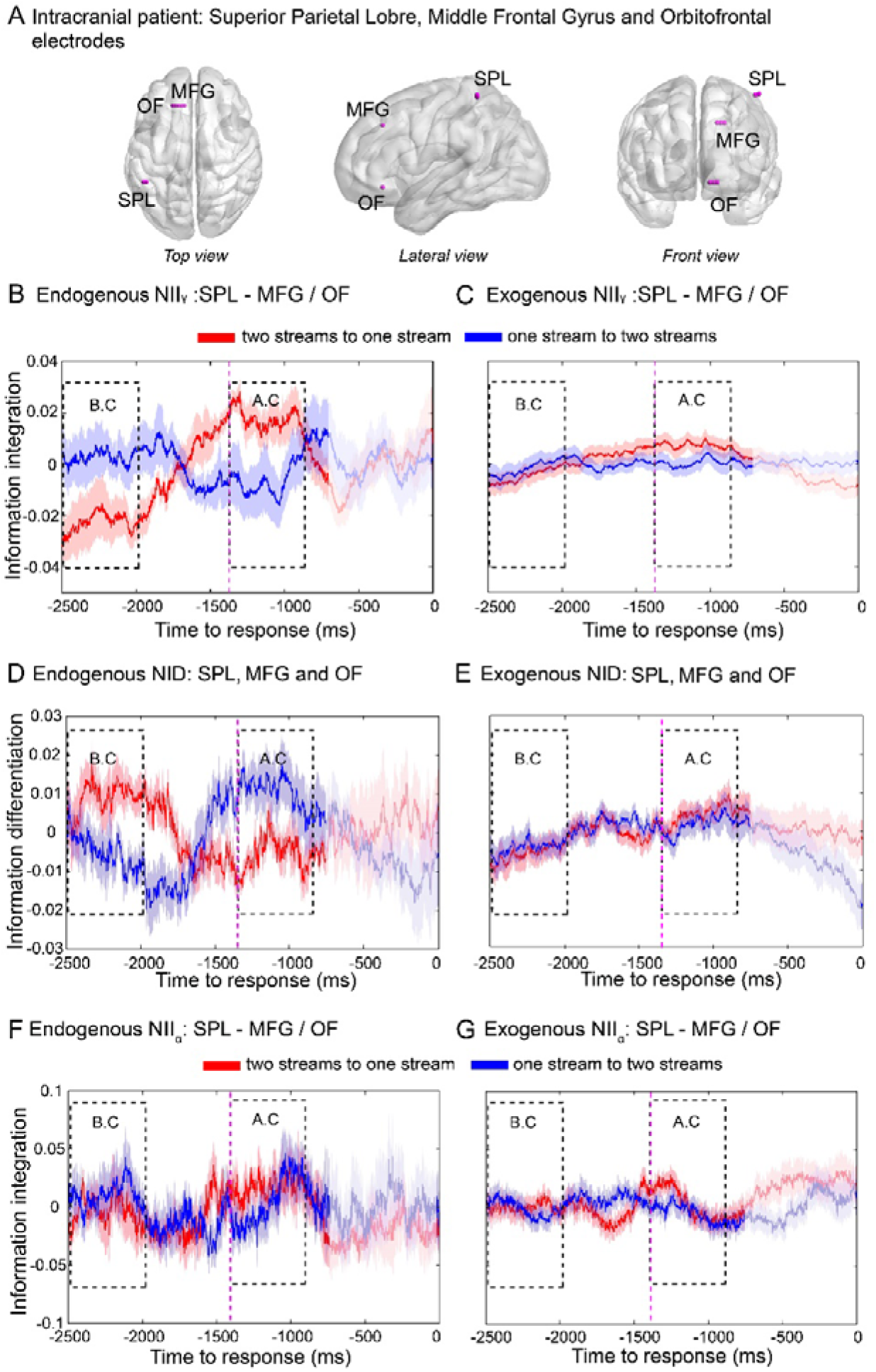
Frontoparietal NII and NID in intracranial recordings. **(a)** Electrodes were implanted in the left superior parietal lobe (SPL), left middle frontal gyrus (MFG) and left orbitofrontal cortex (OF) (Table 1). Representative pair of electrodes (SPL-MFG) showing NII_γ_ and NID of transitions from the two-stream to the one-stream percept (red line) and from the one-stream to the two-stream percept (blue line) in the endogenous **(B, D)** and exogenous **(C, E)** conditions, respectively. **(F, G)** Frontoparietal NII in the alpha band (NOI_α_) in intracranial recordings for the endogenous **(F)** and exogenous condition **(G)**. Purple dashed line marks the mean reaction time (1380 ms) of the exogenous (control) condition of the intracranial data. Statistical analyses were performed as explained in Figure 2. Shaded bars represent s.e.m.

**Figure 5.**
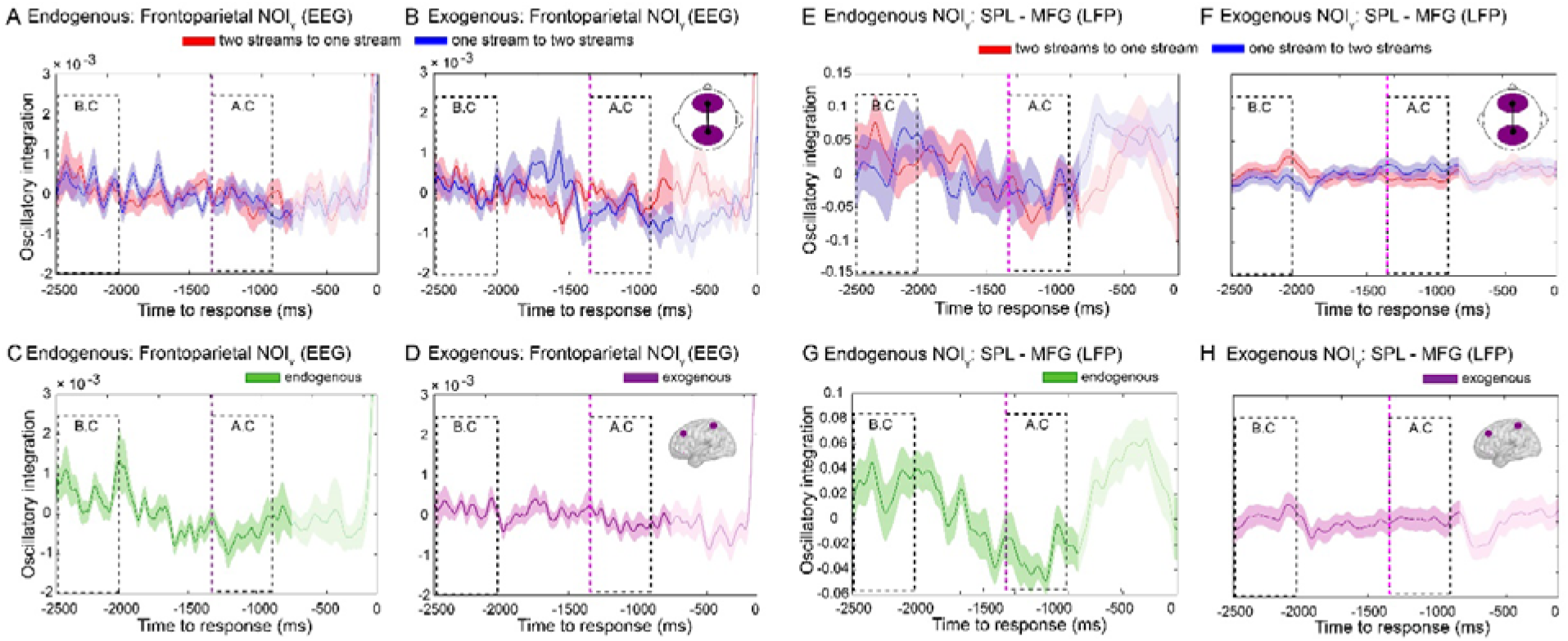
Frontoparietal NOI in the endogenous and exogenous conditions. Dynamics of ongoing NOI_γ_ from the two-stream to the one-stream percept (red line) and from the one-stream to the two-stream percept (blue line) for the endogenous **(A)** and exogenous **(B)** conditions. Ongoing NOI_γ_ during transitions in both directions in the endogenous **(C)** and exogenous **(D)**. Statistical analyses were performed as described in Figure 2. Shaded bars represent s.e.m. **(E-H)** Frontoparietal NOI in the alpha band (NOI_γ_) in intracranial recordings. Ongoing NOI_γ_ during transitions in both directions **(E, F)** in the endogenous **(C)** and exogenous **(D)**. Statistical analyses were performed as described in Figure 2. Error bars represent s.e.m.

### Neural information differentiation (NID) during the endogenous and exogenous conditions

We next investigated the dynamics of neural information differentiation (NID) within frontal and parietal electrodes during bistable perception (Figure 3). The RANOVA revealed a significant triple interaction between condition (endogenous, exogenous), window (before change, after change), and direction (one- to two-stream, two- to one-stream) for NID (*P* = 0.013, 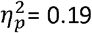, Bf in favour of the alternative = 4.57). Bonferroni’s *post hoc* test revealed higher NID in the two- compared to the one-stream percept in the BC window (*P* = 0.011, 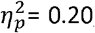, Bf in favour of the alternative = 5.41) (Figure 3A) and a similar pattern in the AC window, showing higher values for two streams than for one stream (*P* = 0.016, Bf in favour of the alternative = 3.88, 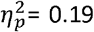) (Figure 3A). These results show that the differentiated percept exhibits higher neurophysiological differentiation than the integrated one.

As was the case for NII_γ_, *post hoc* comparisons showed no NID dependence on percept in the exogenous condition (B.C: *P* = 0.366, Bf in favour of the null = 2.66; A.C: *P* = 0.174, Bf in favour of the null = 1.68) (Figure 3B). Furthermore, no differences were observed in the mean level of NID between endogenous and exogenous conditions (main effect of condition: *F*_1,28_ = 1.39, *P* = 0.248, Bf in favour of the null = 2.12) or between windows (main effect of window: *F*_1,28_ = 0.15, *P* = 0.228, Bf in favour of the null = 3.53), indicating that the overall information differentiation within the frontoparietal network was similar across conditions. Finally, no triple interaction (condition × window × direction) was observed for NID between temporal electrodes (right and left hemispheres) (*F*_1,28_ = 3.59, *P* = 0.069; Bf in favour of the null = 0.86) (Figure 3C, D). In agreement with the NII results, this index of information complexity dissociated endogenous percepts. However, NID showed the opposite pattern compared to NII in terms of the direction of the effects between one and two streams, suggesting that NID is capturing a different but complementary aspect of neural information dynamics associated with conscious percepts.

Consistent with the analysis of NII, we explored NID differences between other electrode regions (i.e. frontofrontal, parietoparietal, temporoparietal and frontotemporal pairs of electrodes). RANOVA revealed no triple interaction (condition × window × direction) between these electrode clusters (Supplementary Figure 6).

**Figure 6.**
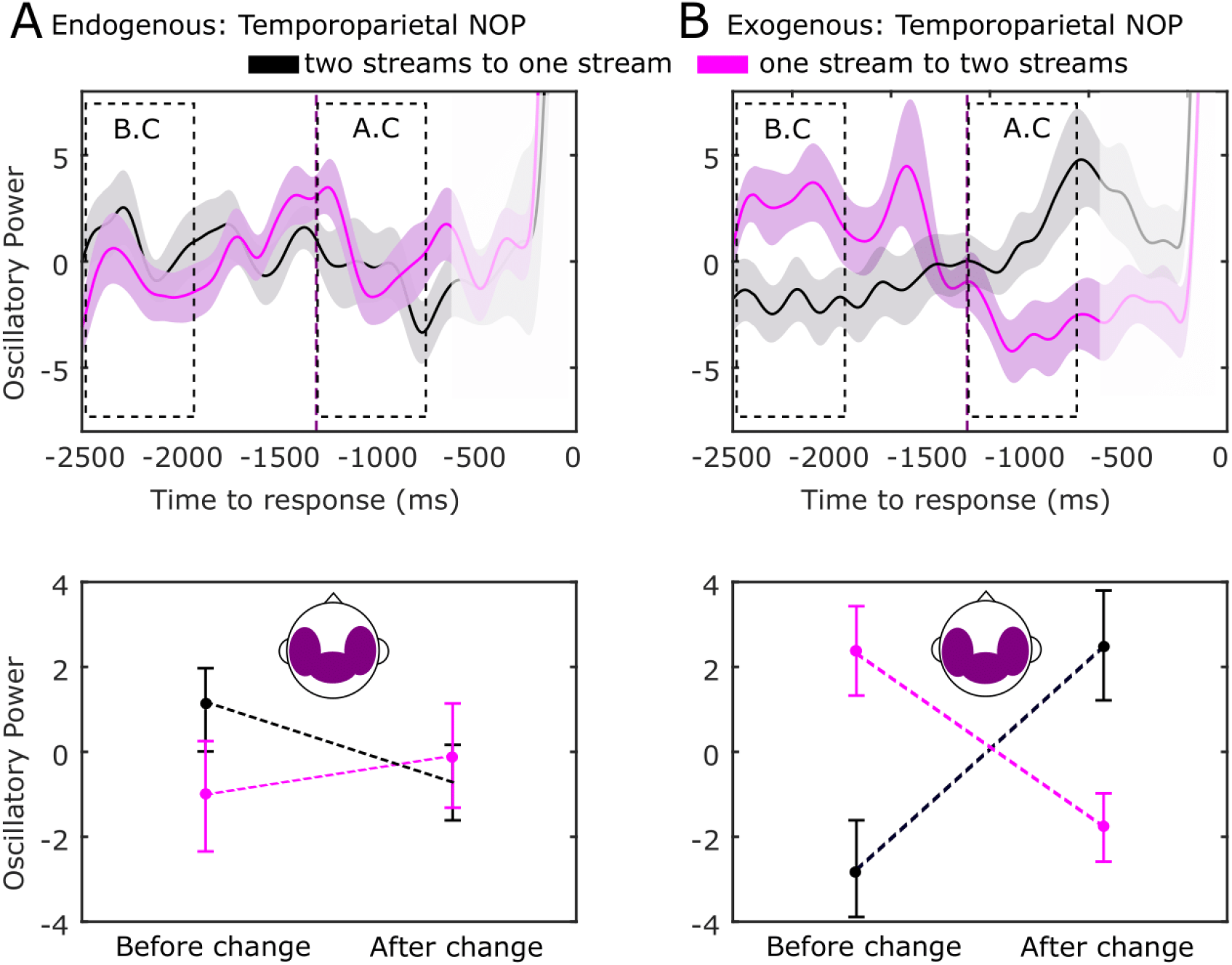
Temporoparietal NOP in the gamma range dissociates alternative exogenous percepts. Neural dynamics of NOP from the two-stream to the one-stream percept (black line) and from the one-stream to the two-stream percept (purple line) for the endogenous **(A)** and exogenous (control) conditions **(B)**. Purple dashed line marks the mean reaction time (1342 ms) of the exogenous (control) condition. Statistical analysis was performed as described in Figure 2. Shaded bars (top row) and error bars (bottom row) represent s.e.m.

### Neural information integration (NII) in human intracortical recordings

In order to validate these findings in a similar manner as elsewhere (Canales-Johnson et al. 2015), we repeated the experiment in a patient implanted with intracranial electrodes for epilepsy surgery. We benefited from the high spatial resolution of intracranial recordings, allowing us to directly test the hypothesis that it is information-sharing specifically between frontal and parietal areas that differentiates between the two auditory percepts. We computed NII_γ_ on intracranial electroencephalography (iEEG) recordings between the superior parietal lobe (SPL) and middle frontal gyrus (MFG), and between SPL and the orbitofrontal cortex (OF) (Table 1) obtained from the intracranial patient performing the same task as above (Figure 4A-C and Supplementary Figure 8). Taking the SPL-MFG and SPL-OF pairs of electrodes together, and as for the healthy participants, the RANOVA showed a triple interaction between condition (endogenous, exogenous), window (before change, after change), and direction (one- to two-stream, two- to one-stream) for NII_γ_ (*F*_1,56_ = 45.19, *P* < 0.001, 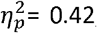, Bf in favour of the alternative > 100). Simple effects within the BC window showed higher NII_γ_ for the one-compared to the two-stream percept in the endogenous (*F*_1,56_ = 44.79, *P* < 0.001, 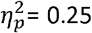, Bf in favour of the alternative > 100) (Figure 4B) but not the exogenous condition (*F*_1,56_ = 0.25, *P* = 0.517, Bf supporting the null = 4.34) (Figure 4C). In the case of the AC window, the one-stream percept again showed higher NII_γ_ than the two-stream percept in the endogenous (*F*_1,56_ = 42.83, *P* < 0.001, 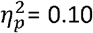, Bf supporting the alternative = 5.10) (Figure 4B) but not in the exogenous condition (*F*_1,56_ = 1.40, *P* = 0.731, Bf in favour of the null = 4.89) (Figure 4C).

In order to further test the frontoparietal specificity of the NII_γ_ modulation, we computed NII_γ_ between SPL and MFG with two unrelated – as per our hypothesis - cortical areas: motor cortex (MC) and posterior insular cortex (PIC) (Supplementary Figure 9). No triple interaction was observed for NII_γ_ between SPL-MC, SPL-PIC, MFG-MC nor MFG-PIC, further confirming the spatial specificity of the intracortical NII_γ_ results.

Furthermore, no differences were found between these percepts in the same iEEG recordings within the alpha band (NII_α_) (*F*_1,56_ = 0.03, *P* = 0.853, Bf in favour of the null = 7.24) (Figure 4F, G). Individual SPL-MFG and SPL-OF pairs for NII_γ_ and NOI_α_ are depicted in Supplementary Figure 8 and Supplementary Figure 10, respectively. Finally, and in agreement with the scalp EEG results, a further iEEG exploratory analysis revealed no triple interaction (RANOVA: condition × window × direction) between SPL-MFG and SPL-OF electrodes in the theta (NII_θ_: Supplementary Figure 11), nor beta (NII_β_: Supplementary Figure 12) bands.

### Neural information differentiation (NID) in human intracortical recordings

Next, we investigated NID dynamics on iEEG signals in the intracranial patient within SPL, MFG and OF (Figure 4D, E). Again, RANOVA showed a triple interaction between (endogenous, exogenous), window (before change, after change), and direction (one- to two-stream, two- to one-stream) (*F*_1,56_ = 24.96, *P* < 0.001, 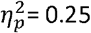, Bf in favour of the alternative = 10.02). In agreement with the scalp EEG results, a simple effects analysis within the BC window showed higher NID for the phenomenologically differentiated percept compared to the phenomenologically integrated percept in the endogenous (*F*_1,56_ = 19.08, *P* < 0.001, 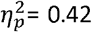, Bf in favour of the alternative > 100) (Figure 4D) but not in the exogenous condition (*F*_1,56_ = 1.92, *P* = 0.171, Bf in favour of the null = 1.94) (Figure 4E). In the case of the AC window, the two-stream percept again showed higher NID than the one-stream percept in the endogenous (*F*_1,56_ = 10.57, *P* = 0.003, 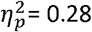, Bf in favour of the alternative = 20.15) (Figure 4D) but not in the control (exogenous) condition (*F*_1,56_ = 0.25, *P* = 0.615, Bf in favour of the null = 4.82) (Figure 7E). These findings demonstrate strong convergent evidence between scalp EEG and direct cortical recordings, further supporting the hypothesis that phenomenology goes along with neurophysiology of conscious percepts, specifically indexed by frontoparietal ongoing activity. Individual SPL-MFG and SPL-OF pairs for NID are depicted in Supplementary Figure 13.

### Neural oscillatory integration (NOI) in the endogenous and exogenous conditions

We next evaluated the theoretical prediction that information dynamics but not oscillatory dynamics of brain activity underpins the emergence of internally-driven percepts (Koch et al. 2016). Thus, we investigated whether neural oscillatory integration (NOI) of ongoing activity might also capture the dynamics of auditory bistability. Specifically, we investigated whether frontoparietal gamma phase synchronization (Weighted Phase-Lag Index (wPLI_γ_)) could differentiate between endogenous percepts. Unlike for NII_γ_, RANOVA revealed no triple interaction between condition (endogenous, exogenous), window (before change, after change), and direction (one- to two-stream, two- to one-stream) for NOI_γ_ (*F*_(1,28)_ = 0.12, *P* = 0.726, Bf in favour of the null = 3.58) (Figure 5A,B). However, a weak interaction between condition (endogenous, exogenous) and window (before change, after change) was found (*F*_(1,28)_ = 4.22, *P* = 0.049, Cohen’s *d* = 0.77, Bf in favour of the alternative = 1.50) (Figure 5C,D). Bonferroni’s *post hoc* test showed that NOI_γ_ significantly decreased in the AC window compared to the BC window in the endogenous (*P* = 0.016, 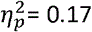, Bf in favour of the alternative = 3.73) (Figure 5C) but probably not in the exogenous condition (*P* = 0.236, Bf in favour of the null = 2.04) (Figure 5D).

Furthermore, the same null result for NOI_γ_ between directions was observed in the intracranial patient (RANOVA (condition × window × direction): *F*_(1,56)_ = 0.12; *P* = 0.726, Bf in favour of the null = 4.73) (Figure 5E,F). As with the scalp EEG, the intracranial patient showed an interaction between condition and window (*F*_(1,56)_ = 13.51, *P* = 0.001, 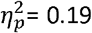, Bf in favour of the alternative = 68.09) in NOI_γ_, showing a decrease in phase synchrony in the AC window compared to the BC window only in the endogenous condition (Bonferroni’s *post hoc* test in endogenous condition: *P* = 0.001, 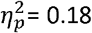, Bf in favour of the alternative = 53.48; and exogenous condition: *P* = 0.147, Bf in favour of the null = 1.92) (Figure 5G,H) between SPL and MFG-OF electrodes. These findings suggest that oscillatory integration does not index the identity of auditory percepts but may relate to percept stability: phase synchrony maxima occur in the BC window (before the end of a stable percept) and phase synchrony minima in the AC window (just at the onset of percept stability).

### Neural oscillatory power (NOP) in the endogenous and exogenous conditions

We finally performed an exploratory analysis aiming at investigating the neural differences during exogenously-driven perception. In order to evaluate the neural response on electrodes sensitive to auditory perturbations, we analysed the neural oscillatory power (NOI) in the same gamma band as previous analyses but focusing on the temporal and parietal electrodes. In the exogenous condition we expected higher gamma spectral power during the integrated (‘ABA-’) as compared to the differentiated (‘AB--’) pattern since the former contains one more tone than the latter, potentially eliciting a greater neural response. RANOVA revealed an interaction between condition (endogenous, exogenous), window (before change, after change) and direction (one- to two-stream, two-to one-stream) for NOP (*F*_(1,28)_ = 4.30, *P* = 0.047, 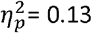, Bf in favour of the alternative = 4.67) (Figure 6). Bonferroni’s *post hoc* test revealed differences in the exogenous condition in line with our prediction. NOP was higher in the one- compared to the two-stream percept in the BC window (*F*_(1,28)_ = 6.37, *P* = 0.001, 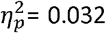, Bf in favour of the alternative = 6.45) (Figure 6) and a similar pattern held in the AC window, with larger values for one stream than for two streams (*P* = 0.001, Bf in favour of the alternative = 7.88, 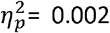) (Figure 6B). No differences in NOP were observed between conscious percepts in the endogenous condition (B.C: *P* = 0.845, Bf in favour of the null = 8.13; *P* = 0.453, Bf in favour of the null = 6.17) (Figure 6A). These results suggest that spectral power within temporal and parietal electrodes is sufficient for indexing conscious percepts when these are externally-driven, that is, when the two patterns (i.e. integrated and differentiated) were physically present in the auditory stimuli.

## DISCUSSION

Here we demonstrate that frontoparietal information dynamics dissociate alternative conscious percepts during internally but not externally driven percepts. By studying the neural dynamics (EEG and direct cortical recordings) of theoretically-motivated information metrics, we show that empirically tractable measures of neural information integration and neural information differentiation map auditory percepts experienced either as perceptually integrated (one-stream) or differentiated (two-stream), respectively. Furthermore, phase synchronization of oscillatory gamma activity in the frontoparietal network does not differentiate between auditory percepts, nor does the information between temporal networks. Finally, when perception was driven externally (no bistability, control condition) by a change in the sound stream, neural oscillatory power distinguished between percepts but information measures did not.

### Neural correlates of consciousness and the instantiation of meaningful signatures of information integration and differentiation

Our results expand the understanding of the neural correlates of consciousness (NCC) (Koch et al. 2016) in several ways. First, our experimental findings directly support information-based theories of consciousness (Dehaene and Changeux 2011; Dehaene et al. 2014; Tononi et al. 2016); in their current instantiation, the Integrated Information Theory (IIT) (Tononi et al. 2016) and the Global Neuronal Workspace Theory (GNWT) (Dehaene and Changeux 2011) of consciousness both emphasize the role of information exchange in generating conscious percepts. Although both theories conceptualize information differently, their proposed empirical indices are based on the classical Shannon-entropy information framework. Using these measures of information dynamics, our results show convergent evidence supporting both GNWT and IIT predictions by demonstrating a role of neural information in the emergence of contents of consciousness. However, the interpretation of these results in light of the IIT and GNWT is somewhat different.

The GNWT provides a framework for evaluating the process of becoming aware of a percept (i.e. the conscious access for a stimulus), specifically in our case the experience of perceiving one or two auditory streams. Of the proposed key features of neural processing that the GNWT supports is the modulation of frontoparietal networks when gaining conscious awareness. This is why we set out to test the frontoparietal network at the electrode level (EEG) and the intracranial cortical level (iEEG).

In its original conception, the GNWT did not specifically proposed integration measures that would be neural signatures of gain of conscious awareness. Later on, however, in a landmark paper Gaillard et al in 2009 showed connectivity and granger-causality calculations that signalled the access to consciousness in a masking paradigm in intracranial patients (Gaillard et al., 2009). Echoing the need to better define what a putative measure of brain integration is, we searched for those measures that could be used to assess integration information and differentiation in the time dynamics of the stream of consciousness of the current task.

Although most of IIT’s empirical work has been focused on finding proxy measures for distinguishing levels of consciousness (e.g. wakefulness versus sleep), recent efforts have been put in investigating perceptual integration and differentiation at the first-person, phenomenological level using time-resolved experimental paradigms (Boly at al., 2015). We think that this current framework allows for testing the concepts of integration and differentiation in the context of the study of the first-person, content of consciousness (i.e. hearing one-stream vs two-stream stimuli) rather than on the study of the third-person, state of consciousness (e.g. being awake vs being asleep). We believe our study substantially improves this approach for three main reasons. First, the repertoire of conscious contents is controlled (i.e. one vs two auditory streams). Second, the physical stimulation remains constant while perception does not. Finally, our EEG/iEEG information metrics possess higher temporal resolution than those used in fMRI studies, allowing the finely-grained characterization of the phenomenology of conscious contents.

It is in this respect that we think that concepts taken from both theoretical frameworks (GNWT and IIT) are useful to explain some cognitive process that we believe are at play in bistable perception. We do not claim that this unifies the theories, but that it is possible to make use of parts of them to frame results that can help understanding the neural signatures of contents of consciousness. An important caveat of this conceptualization is that the theoretical argumentations from GNWT and IIT do not make explicit predictions about the possible differences between internally and externally driven contents of consciousness, while we explicitly test this difference.

Our results suggest a differential role of information integration vs. information differentiation in the emergence of conscious percepts from those proposed before. According to IIT, the neural activity associated with conscious percepts should reflect the joint presence of neurophysiological integration and neurophysiological differentiation. Under this theoretical framework, integration is expected – in principle – to be paralleled by differentiation of neural activity. Contrary to this prediction, our results show dissociation between neurophysiological integration and neurophysiological differentiation of frontoparietal ongoing activity. Interestingly, while the phenomenologically integrated percept (one-stream) showed a relative increase in NII and relative decrease in NID, the perceptually differentiated percept (two-stream) exhibited the opposite pattern, that is, a decrease in NII and increase in NID. Together, these dissociated patterns suggest that each measure is instead directly associated with the phenomenology of internally-driven conscious percepts: whereas information integration of neural activity is capturing phenomenological integration (one-stream percept), information differentiation may be capturing phenomenological differentiation (two-stream percept).

### Neural signatures of the emergence of internally- and externally-driven conscious contents

We have demonstrated a potential mechanistic role of information integration and differentiation in the formation of endogenous percepts. Why do information metrics capture the neural dynamics of internally-driven conscious percepts? Coordination in the brain has been classically studied by computing phase synchronization between neural oscillations (Uhlhaas et al. 2009; Engel et al. 2013). Synchronization is a highly-ordered form of neural coordination that primarily captures the linear (or proportional) phase relationship between signals at specific frequencies (phase-locking). Thus, a mechanism of coordination-by-synchrony captures only certain regimes of neural coordination that are periodic. However, brain dynamics exhibit both a tendency to integrate information (synchronization) and a tendency for the components to differentiate information (independent function) (Dehaene and Changeux 2011; Tognoli and Kelso 2014; Tononi et al. 2016). During auditory bistability, conscious percepts typically alternate continuously without becoming locked into any one percept for long periods. We propose that underlying this dynamical process are ensembles of neurons that are repeatedly assembled and disassembled, and that this non-trivial dynamic might be instantiated by a mechanism of coding-by-information that captures complex, nonlinear patterns of neural activity and not merely simple proportional associations between neural signals. In fact, we have recently shown that wSMI (NII) performs better at capturing highly non-linear interactions than wPLI (NOI) between long-distance neural signals by using realistic brain simulations (Imperatori et al., 2019).

Contrasting the emergence of internally-driven conscious percepts, the two auditory sequences presented during the exogenous conditions were distinguished by a difference in spectral power in the gamma band. The difference was observed mainly in the temporoparietal electrodes and the direction of the effect is consistent with the idea that an extra tone in the sequence elicited a higher neural response, i.e. that the ABA- sequence showed higher gamma power than the AB-- sequence. Thus, contrary to the information-theory metrics, we suggest that the spectral gamma power response is linearly (or proportionally) related to the physical characteristics of the stimulus itself (exogenous condition), and not to the internal interpretation of the bistable sequence, as no differences were observed in the endogenous condition. Hence for the exogenous condition, while the report is the same (one or two streams), the neural signatures point a different process in the brain when the nature of the contents differs.

### Conscious auditory percepts and ongoing neural activity

The frontoparietal patterns of neural information are a manifestation of the interaction between the external stimulation and endogenous, ongoing brain activity, as opposed to activation purely imposed by the auditory stimuli. Thus, NII and NID patterns do not merely reflect stimulus-driven neural activity but rather the intrinsic coordination of endogenous frontoparietal neural activity. Indeed, in the endogenous condition of our study, internally generated changes in neural activity were associated with changes in conscious percepts in the complete absence of any change in the auditory stimuli. In line with our results, recent studies in the visual system have shown that long-distance integration of ongoing oscillations reflects internally coordinated activity associated with conscious perception (Hipp et al. 2012; Engel et al. 2013; Helfrich et al. 2016). Here, by directly measuring the amount of information integration and information differentiation contained in the ongoing neural activity, we demonstrate a functional role of information dynamics in the emergence of auditory conscious percepts. Our approach provides a framework for examining whether this relationship between neural and perceptual integration and differentiation generalizes, for example to the perception of ambiguous stimuli in other sensory modalities, or across modalities.

Although the pioneering electrophysiological studies supporting the active role of ongoing activity in perception and cognition date from the 70’s (Freeman 1976, 2000), over the last decade ongoing brain activity has been mainly studied in the context of “resting state networks” (Fox and Raichle 2007). In these recent studies, fluctuations in ongoing activity between spatially segregated networks (brain regions) are correlated when a participant is not performing an explicitly defined task. In the present study, by taking advantage of the high temporal resolution of EEG and direct cortical recordings, we show that patterns of neural information are transiently coordinated during the active discrimination of internally generated auditory percepts. Furthermore, our results also allowed us to differentiate the contribution of information in frequency-space, showing that gamma but not alpha NII differentiates auditory percepts during bistable perception.

### Auditory bistable perception and frontoparietal activity

Our results also suggest a fruitful new approach to conceptualizing and investigating bistable auditory perception. We demonstrate that the dynamics of auditory endogenous percepts can be associated with long-distance coordination of neural activity. For the ABA- patterns presented here, while both one-stream and two-stream percepts depend on the integration of tones over time, the latter also requires integration across tone frequency. We argue that this additional integration may draw on, or be reflected in, an increase in information sharing in ongoing frontoparietal neural activity. Additionally, maintenance of two distinct perceptual streams of sounds was associated with more differentiated neural patterns than when a single stream was perceived. How these findings relate to previous demonstrations of greater stimulus-locked activity in auditory cortex (Gutschalk et al. 2005; Hill et al. 2011; Szalardy et al. 2013; Billig et al. 2018; Sanders et al. 2018) and greater BOLD response in intraparietal sulcus (Cusack 2005; Hill et al. 2011) for two streams is presently unclear. However, the involvement of neural circuits extending beyond sensory regions in parsing the auditory environment is not surprising given that attention switching (Billig and Carlyon 2016), linguistic knowledge (Billig et al. 2013), and predictability (Winkler and Schröger 2015) have been shown to affect auditory streaming.

These findings provide convergent evidence about the role of frontoparietal networks in the dynamics of formation and maintenance of internally-driven conscious contents. Research in visual bistability has focused on characterizing content-related activity predominantly in local brain areas or networks (Sterzer et al. 2009). Of those few studies that have expanded their scope to associative cortices and wider networks, one has recently proposed mechanistic accounts on visual percepts using multivariate pattern analysis (MVPA) of fMRI data (Wang et al. 2013). The results showed differential patterns of BOLD activity in high-order frontoparietal regions between visual percepts during bistable perception. The present study complements these results by showing a role for the frontoparietal network in indexing percepts in the auditory modality. Furthermore, the temporal resolution of our EEG and iEEG data enabled us to characterize the fine-grained temporal dynamics of neural information integration associated with specific auditory percepts within the frontoparietal network. These results represent convergent evidence towards a possible general mechanism of information integration underlying the emergence of the contents of consciousness under invariant stimulation.

In conclusion, we have presented experimental evidence that conscious percepts may be supported by different neural mechanisms depending on whether they are internally or externally driven. We have highlighted the stark contrast between fleeting endogenous percepts, where neural integration and differentiation parallel the corresponding integrated and differentiated percepts, and those that are externally triggered, for which no differences between information-theory measures were observed. However, when conscious percepts are driven by changes in the external stimulation, fluctuations in spectral power are sufficient for indexing perceptual integration and differentiation.

Importantly, the conceptual mapping between phenomenology and the neurophysiology that we have highlighted here should be considered as a fruitful approach for measuring the different dimensions of phenomenology in an experimentally testable manner. In light of some of the main current theoretical frameworks of conscious perception -IIT and GNWT- we provide additional experimental scaffolding to further understand what we have in mind at any given time.

## Acknowledgments

We thank Robert Carlyon, Simon van Gaal, Jean-Phillippe Lachaux, Anat Arzi, William J. Harrison, Daniel Bor and Victor Lamme for contributing to valuable discussion and insights. This manuscript is dedicated to the memory of Prof. Walter J. Freeman (1927 - 2016) whose pioneering work on Neurodynamics has inspired and ignited countless meaningful insights during the execution of this project.

## Notes

**Funding** This research was supported by a Wellcome Trust Biomedical Research Fellowship WT093811MA and the Chilean National Fund for Scientific and Technological Development Grant 1171200.

**Conflict of Interest** None declared.

