## Supplementary Figures for "Dissociable neural information dynamics of perceptual integration and differentiation during bistable perception"

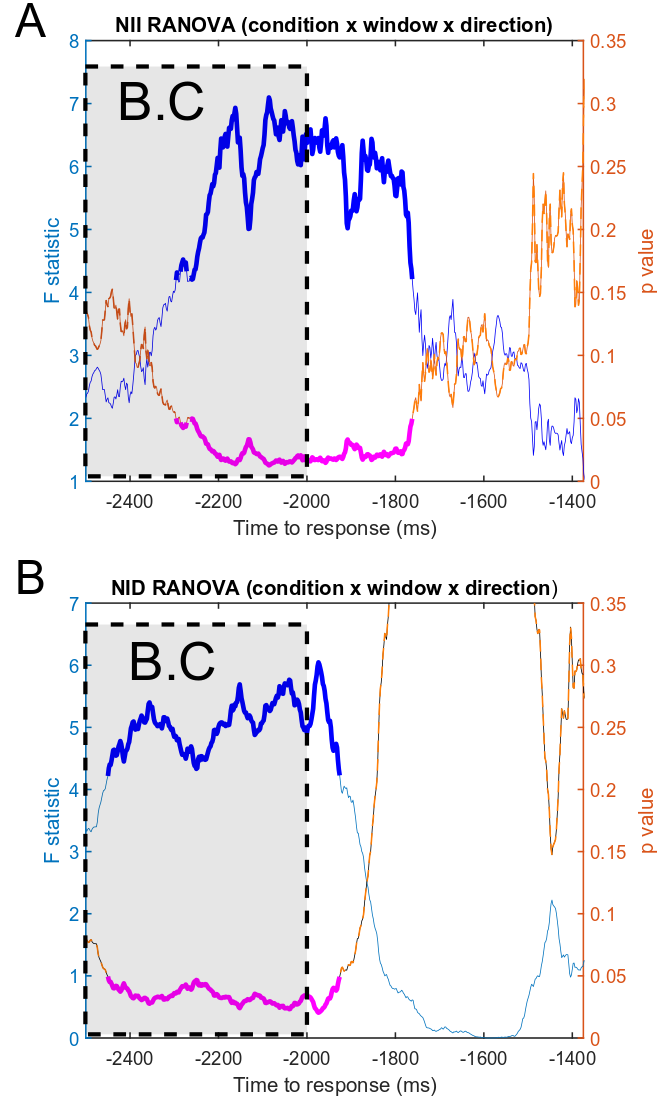

**Supplementary Figure 1. Sliding-window statistical analysis for NII and NID of Figure 2 and Figure 3.** In order to show the statistical effects as a function of the window size, we ran a sliding window RANOVA for both the NII **(A)** and NID **(B)** effects reported in Figures 2 and 3, respectively. RANOVA factors are described in the Materials and Methods, i.e. 3 within-participant factors: condition (endogenous, exogenous), window (before-change (B.C.), after-change (A.C)), and direction (one- to two-stream, two- to one-stream). We first computed the mean NII and NID values in the AC window for the endogenous and exogenous conditions (500 ms; from -1342 to -842 ms). RANOVA was then computed using the fixed mean AC window value described above, and a variable value of the BC window for each RANOVA test (1 point per comparison) in a sliding-window manner from -2500 to -1342 ms. Following this procedure, we obtained an F-statistic (blue line) of the triple interaction (condition x window x direction) and its corresponding P value (orange line) for each time point during the entire period preceding the mean reaction time of the exogenous condition (1342 ms). We concluded that the optimal time-window size, i.e. when most of the time points are significant (F-stats values with a P < 0.05 depicted in dark blue and pink, respectively) were not exactly within the time-windows used for our hypothesis-driven NII and NID calculations in the main manuscript (B.C: -2500 to -2000 ms; shaded rectangle in discontinuous lines).

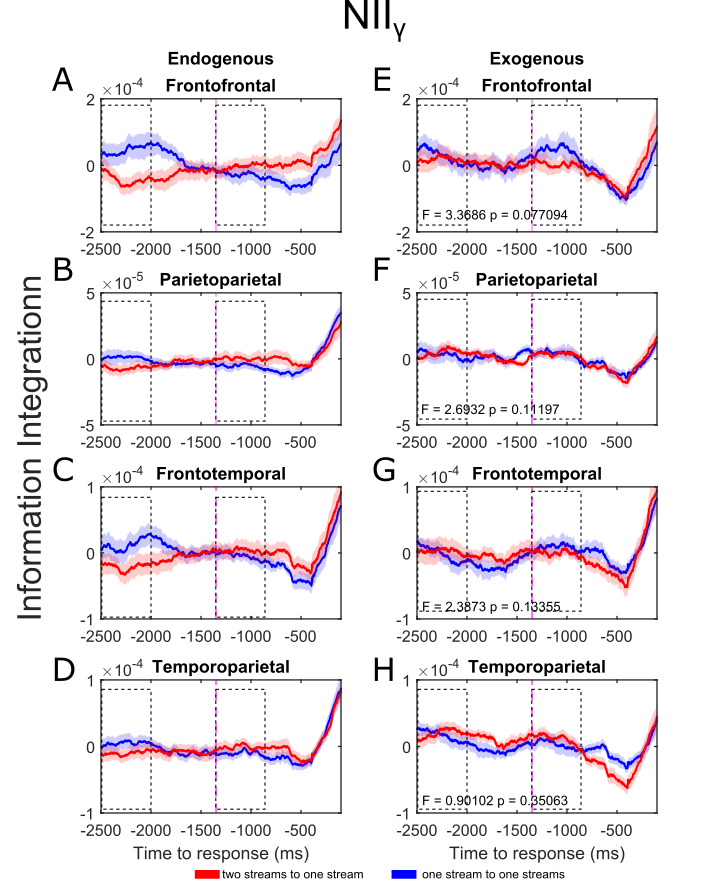

**Supplementary Figure 2. Neural information integration in the gamma range (NII_γ_) between distributed electrode pairs.** Dynamics of NII from the two-stream to the one-stream percept (red line) and from the one-stream to the two-stream percept (blue line) for the endogenous **(A, B, C, D)** and exogenous (control) conditions **(E, F, G, H)**. NID dynamics for the endogenous and exogenous (control) conditions for Frontofrontal **(A, E)**, Parietoparietal **(B, F)**, Frontotemporal **(C, G)** and Temporoparietal **(D, H)** electrode pairs. Purple dashed line marks the mean reaction time (1342 ms) of the exogenous (control) condition. Statistical analysis was performed as described in Figure 2. Shaded bars (top row) and error bars (bottom row) represent s.e.m.

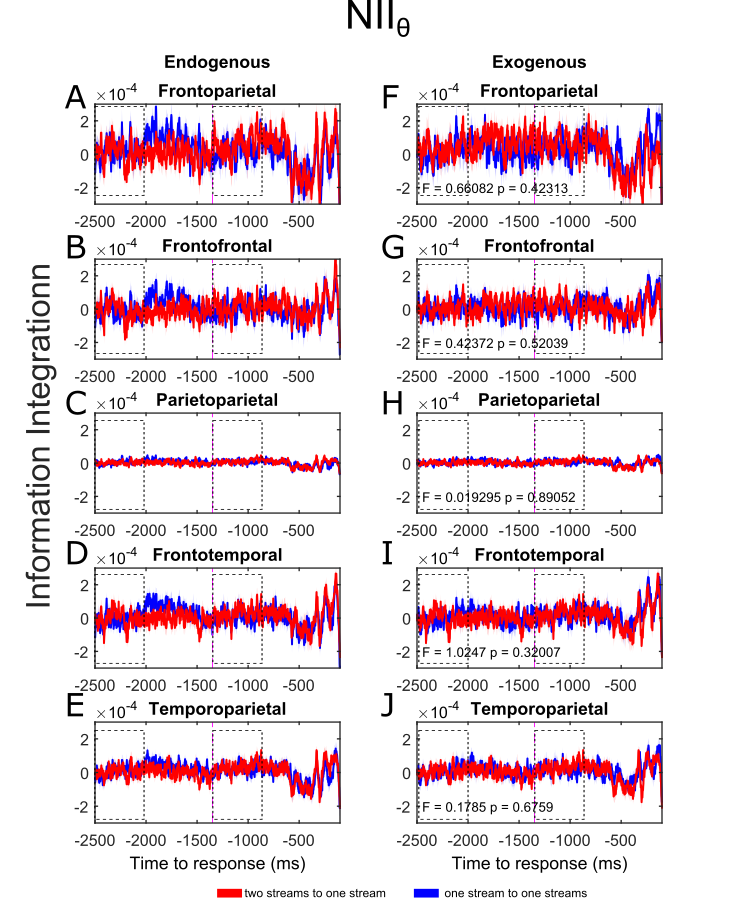

**Supplementary Figure 3. Neural information integration in the theta range (NII_θ_) between distributed electrode pairs.** Neural dynamics of NII from the two-stream to the one-stream percept (red line) and from the one-stream to the two-stream percept (blue line) for the endogenous **(A, B, C, D, E)** and exogenous (control) conditions **(F, G, H, I, J)**. NID dynamics for the endogenous and exogenous (control) conditions for Frontoparietal **(A, F)**, Frontofrontal **(B, G)**, Parietoparietal **(C, H)**, Frontotemporal **(D, I)** and Temporoparietal **(E, J)** electrode pairs. Purple dashed line marks the mean reaction time (1342 ms) of the exogenous (control) condition. Statistical analysis was performed as described in Figure 2. Shaded bars (top row) and error bars (bottom row) represent s.e.m.

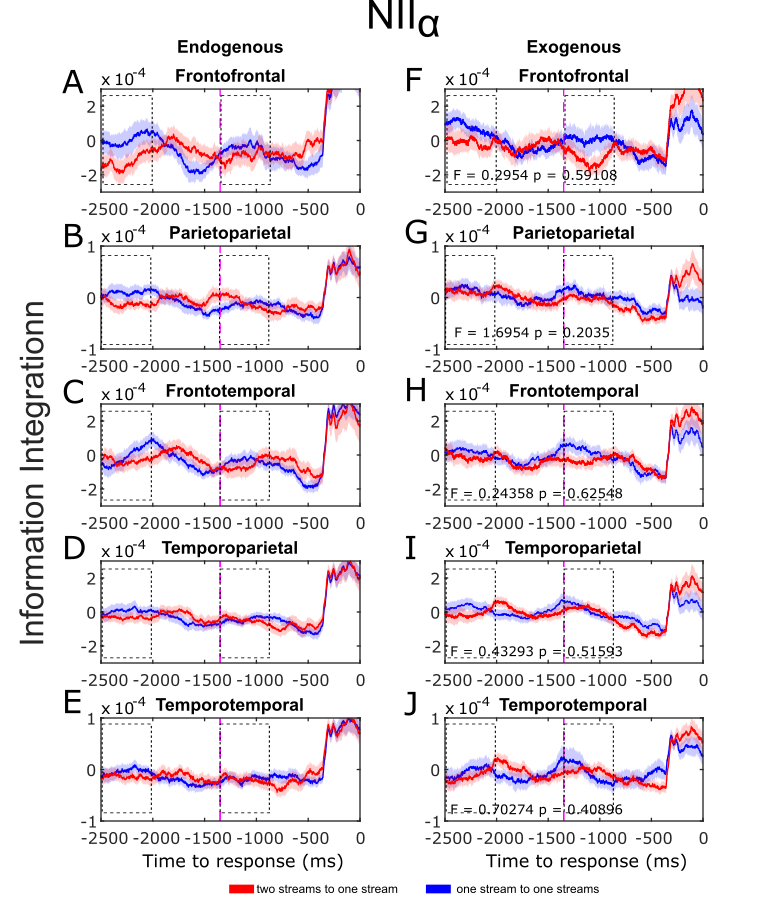

**Supplementary Figure 4. Neural information integration in the alpha range (NII_α_) between distributed electrode pairs.** Neural dynamics of NII from the two-stream to the one-stream percept (red line) and from the one-stream to the two-stream percept (blue line) for the endogenous **(A, B, C, D, E)** and exogenous (control) conditions **(F, G, H, I, J)**. NID dynamics for the endogenous and exogenous (control) conditions for Frontofrontal **(A, F)**, Parietoparietal **(B, G)**, Frontotemporal **(C, H)**, Temporoparietal **(D, I)**, and Temporotemporal **(E, J)** electrode pairs. Purple dashed line marks the mean reaction time (1342 ms) of the exogenous (control) condition. Statistical analysis was performed as described in Figure 2. Shaded bars (top row) and error bars (bottom row) represent s.e.m.

**Supplementary Figure 5. Neural information integration in the beta range (NII_β_) between distributed electrode pairs.** Neural dynamics of NII from the two-stream to the one-stream percept (red line) and from the one-stream to the two-stream percept (blue line) for the endogenous **(A, B, C, D, E)** and exogenous (control) conditions **(F, G, H, I, J)**. NID dynamics for the endogenous and exogenous (control) conditions for Frontoparietal **(A, F)**, Frontofrontal **(B, G)**, Parietoparietal **(C, H)**, Frontotemporal **(D, I)** and Temporoparietal **(E, J)** electrode pairs. Purple dashed line marks the mean reaction time (1342 ms) of the exogenous (control) condition. Statistical analysis was performed as described in Figure 2. Shaded bars (top row) and error bars (bottom row) represent s.e.m.
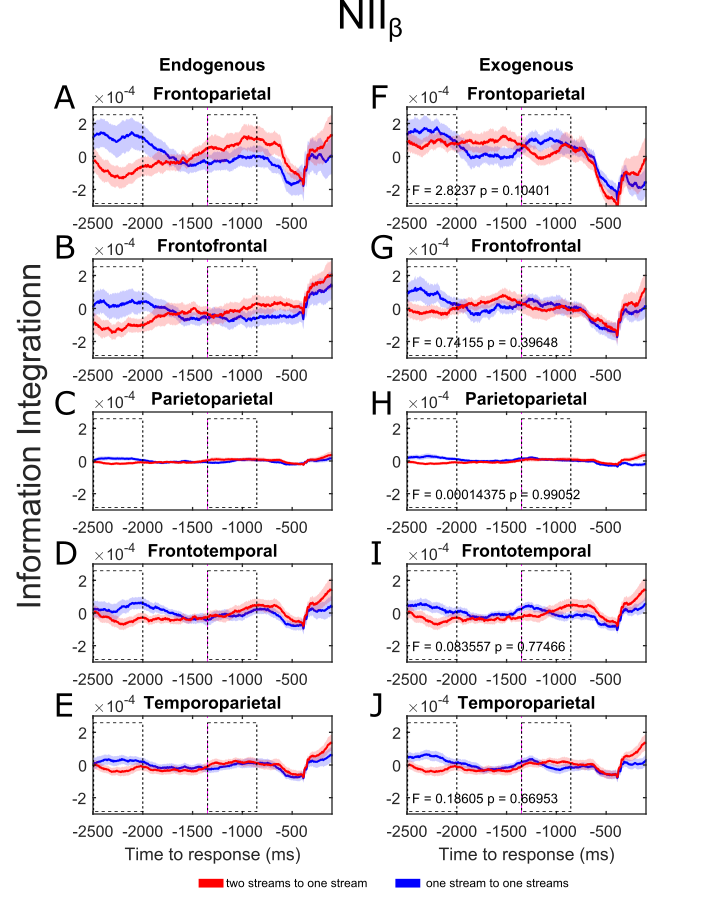

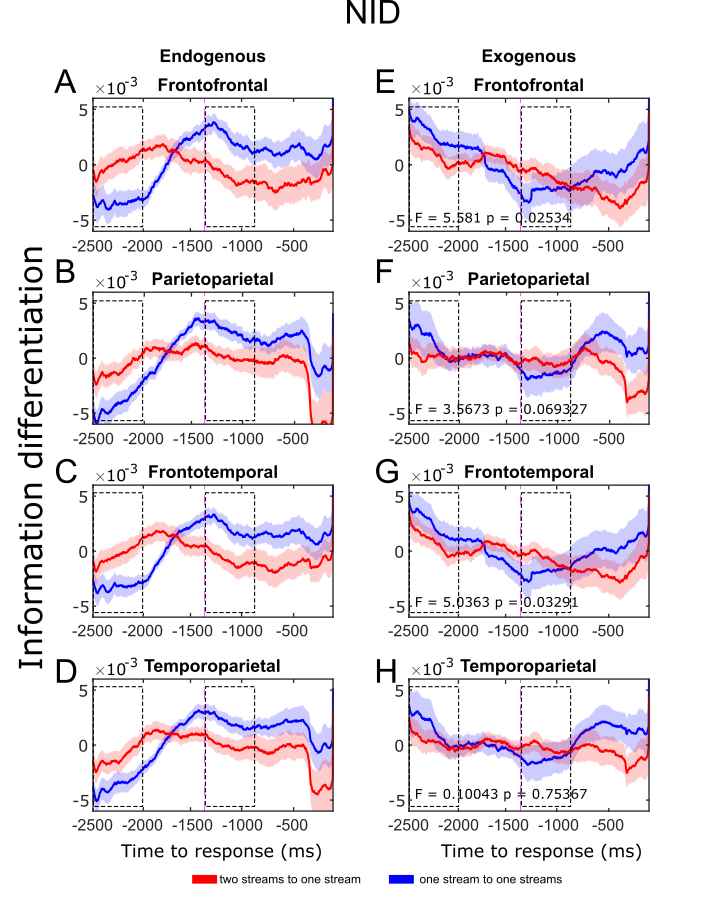

**Supplementary Figure 6. Neural information differentiation (NID) between distributed electrode pairs.** Neural dynamics of NID from the two-stream to the one-stream percept (red line) and from the one-stream to the two-stream percept (blue line) for the endogenous **(A, B, C, D)** and exogenous (control) conditions **(E, F, G, H)**. NID dynamics for the endogenous and exogenous (control) conditions for Frontofrontal **(A, E)**, Parietoparietal **(B, F)**, Frontotemporal **(C, G)** and Temporoparietal **(D, H)** electrode pairs. Purple dashed line marks the mean reaction time (1342 ms) of the exogenous (control) condition. Statistical analysis was performed as described in Figure 2. Shaded bars (top row) and error bars (bottom row) represent s.e.m.

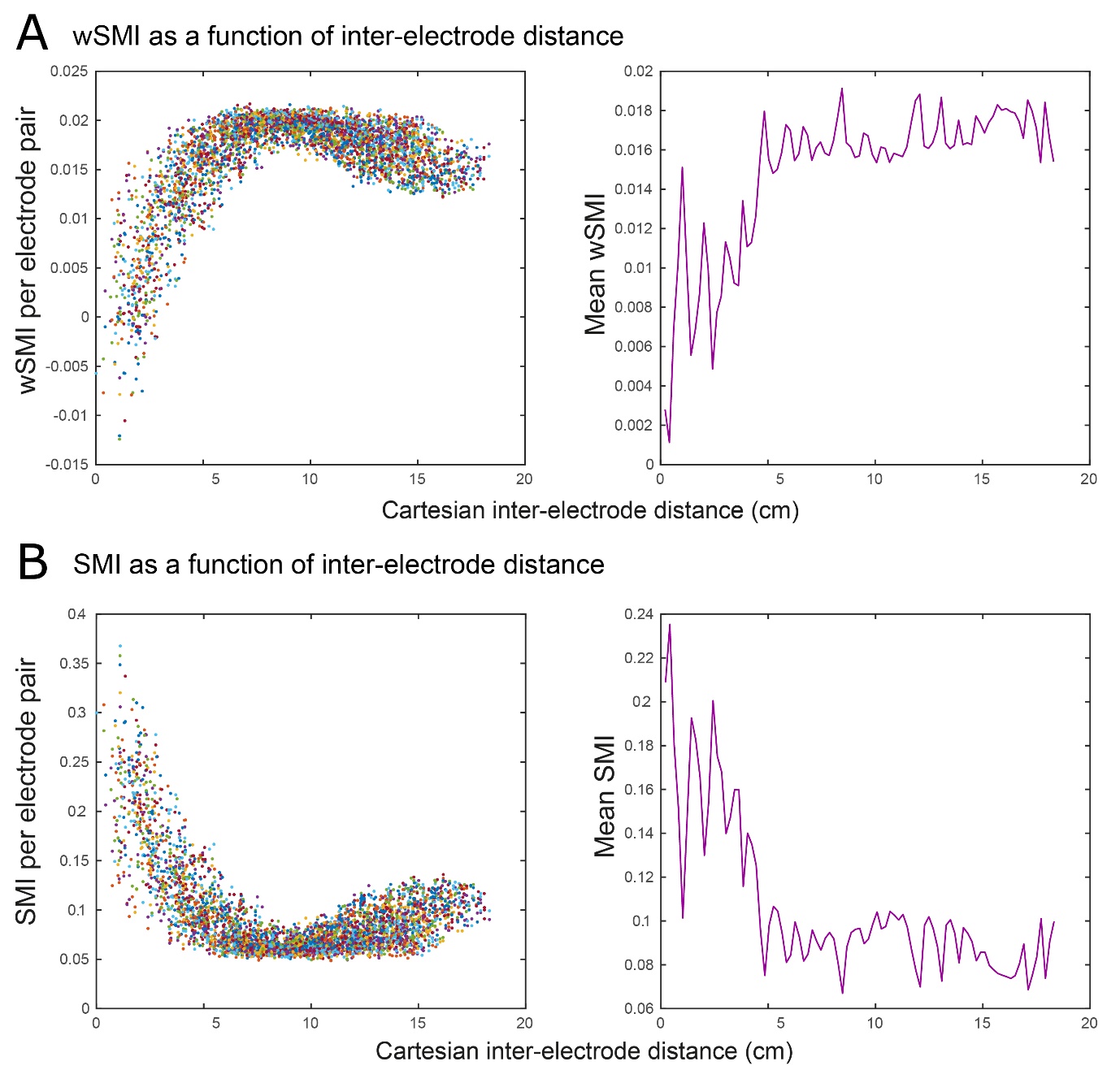

**Supplementary Figure 7. SMI and wSMI variation with inter-electrode distance. (A)** individual wSMI pairs (left panel) and mean wSMI (right panel) across participants as a function of inter-electrode distance in the gamma range. For distances below 5 cm, wSMI quickly dropped toward zero, as expected given that the weighted version of this measure was designed to eliminate common source artifacts (King et al., 2013; Sitt et al., 2014). **(B)** Individual SMI (left panel) and mean SMI (right panel) cross participants as a function of inter-electrode distance in the gamma range. On the contrary, the non-weighted version of the metric (SMI) exhibited the highest values below 5 cm, typically produced by common source artifacts (King et al., 2013; Sitt et al., 2014). wSMI and SMI values were calculated on the AC and BC windows across participants and plotted altogether.

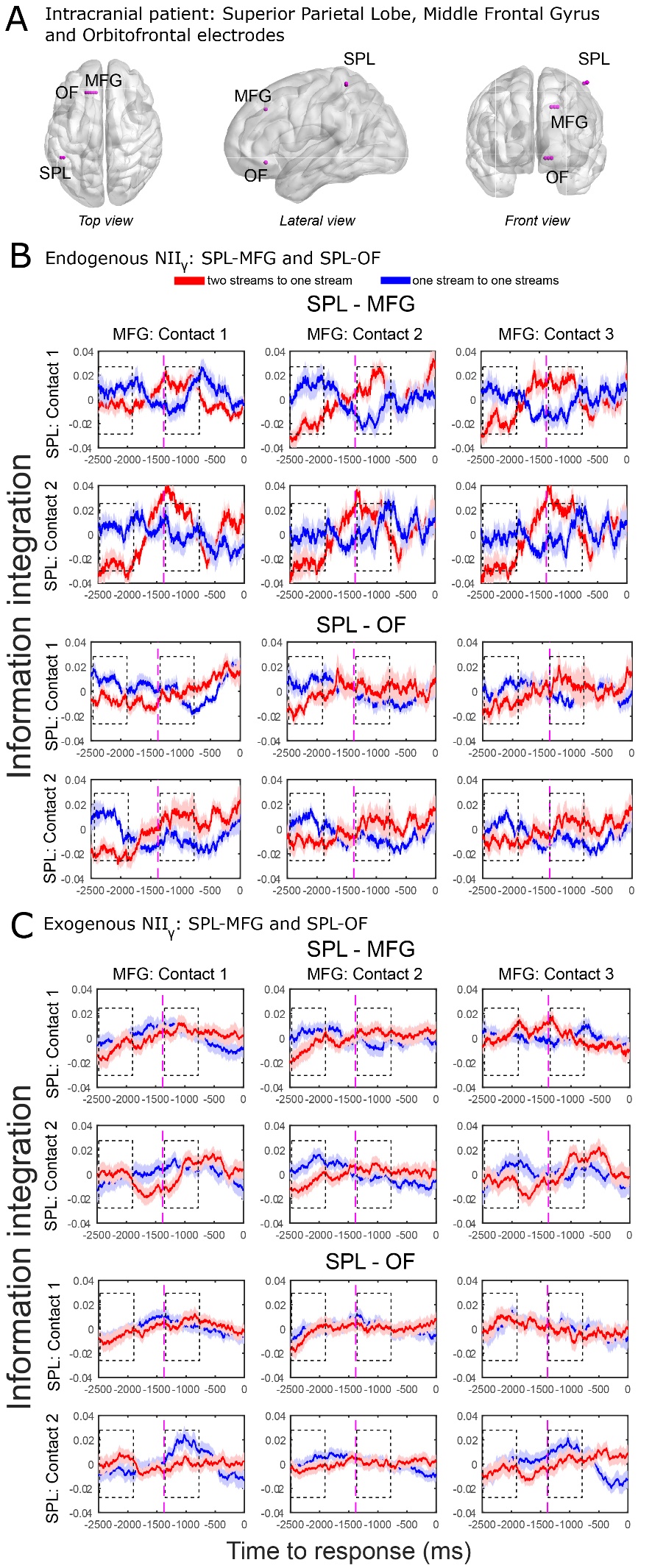

**Supplementary Figure 8. iEEG neural information integration in the gamma range (NII_γ_).** **(A)** Electrodes were implanted in the left superior parietal lobe (SPL: 2 contacts), left middle frontal gyrus (MFG: 3 contacts) and left orbitofrontal cortex (OF: 3 contacts) (Table 1). SPL-MFG and SPL-OF pairs showing NII_γ_ of transitions from the two-stream to the one-stream percept (red line) and from the one-stream to the two-stream percept (blue line) in the endogenous **(B)** and exogenous **(C)** conditions. Purple dashed line marks the mean reaction time (1380 ms) of the exogenous (control) condition of the intracranial data. Statistical results are described in the main text.

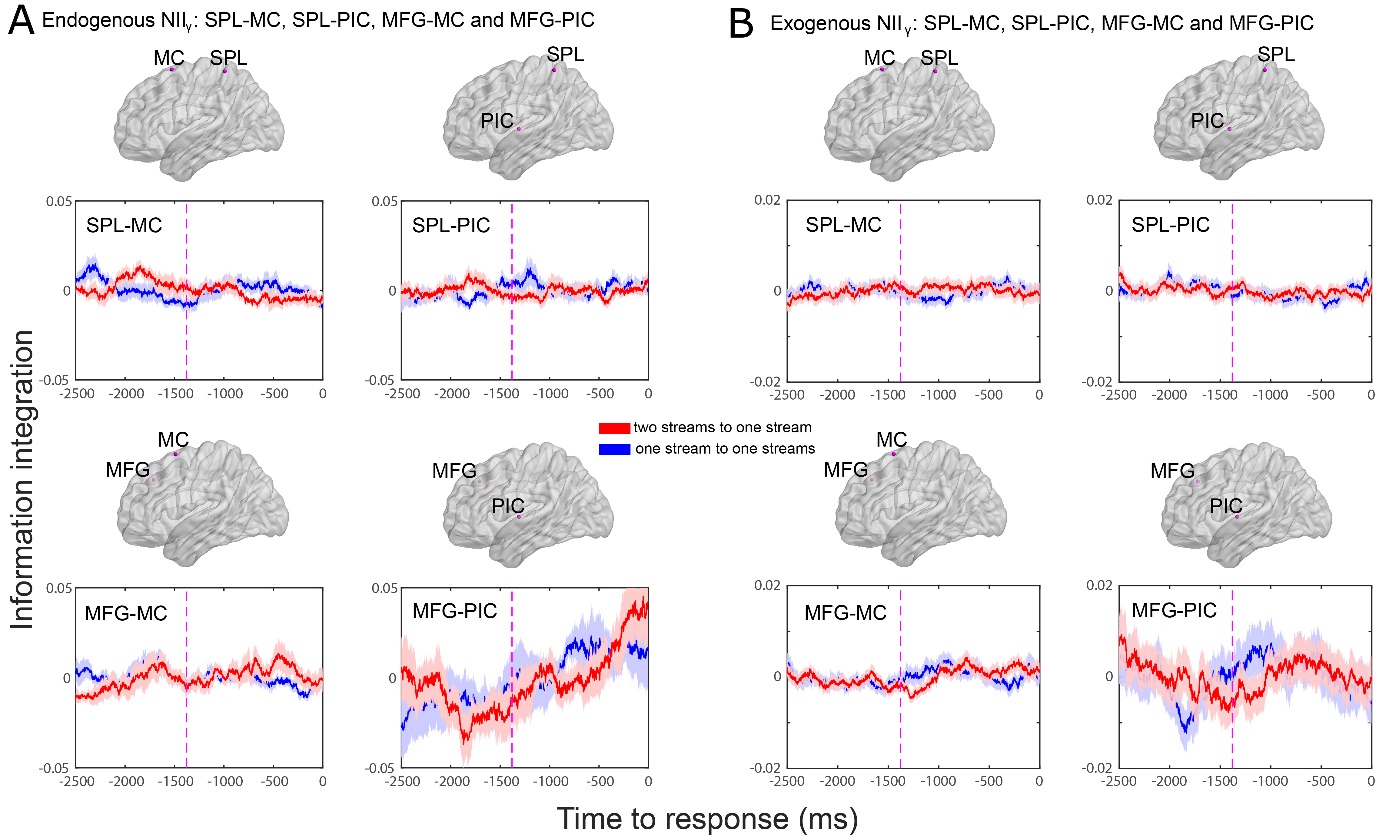

**Supplementary Figure 9. iEEG neural information integration in the gamma range (NII_γ_) for control electrode pairs (A, B)** Electrodes were implanted in the left superior parietal lobe (SPL), left middle frontal gyrus (MFG), left motor cortex (MC) and left posterior insular cortex (PIC) (Table 1). SPL-MC, SPL-PIC, MFG-MC and MFG-PIC pairs show NII_γ_ of transitions from the two-stream to the one-stream percept (red line) and from the one-stream to the two-stream percept (blue line) in the endogenous **(B)** and exogenous **(C)** conditions. Purple dashed line marks the mean reaction time (1380 ms) of the exogenous (control) condition of the intracranial data. No differences were observed for these pairs of electrodes. No triple interaction was found between condition x window x direction (SPL-MC: *F*_1,56_ = 0.805; *P* = 0.373, SPL-PIC: *F*_1,56_ = 0.380; *P* = 0.540, MFG-MC: *F*_1,56_ = 0.276; *P* = 0.601, MFG-PIC: *F*_1,56_ = 0.361; *P* = 0.551)

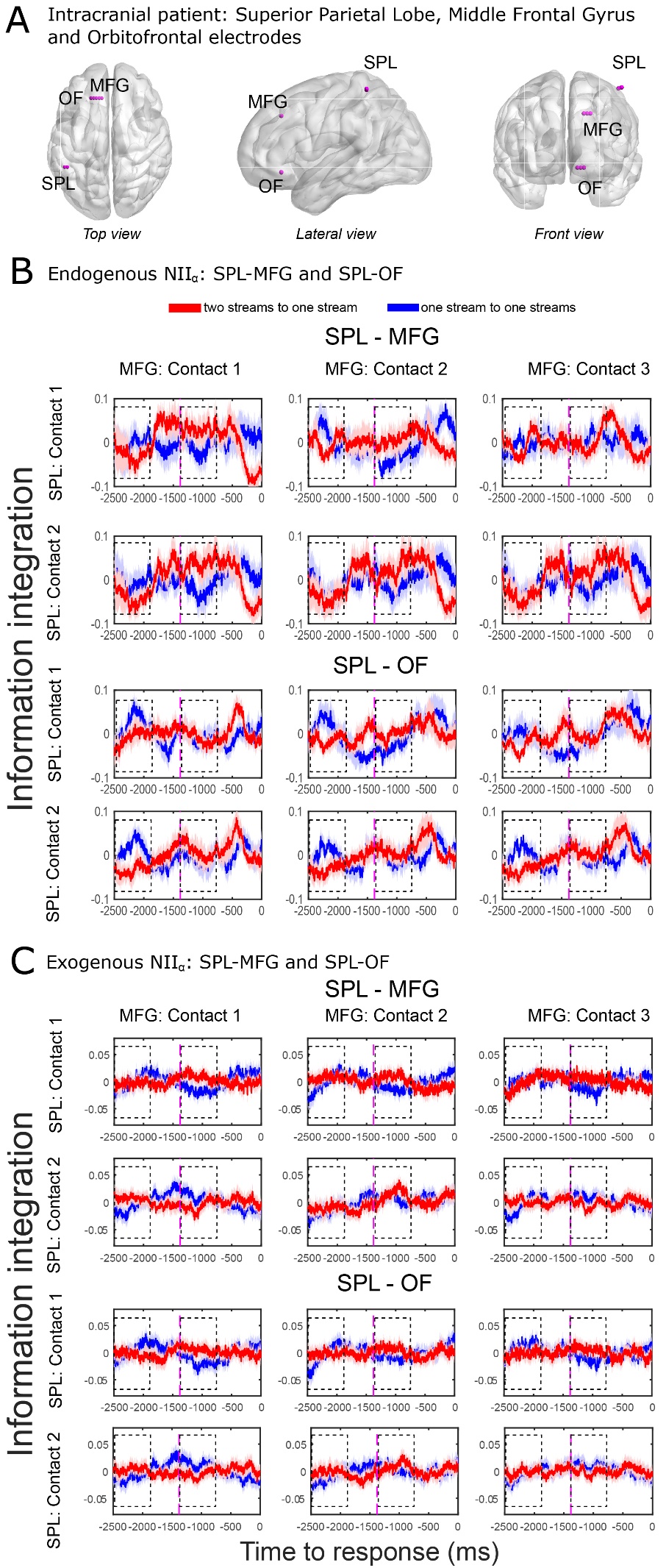

**Supplementary Figure 10. iEEG neural information integration in the alpha range (NII_α_).** **(A)** Electrodes were implanted in the left superior parietal lobe (SPL: 2 contacts), left middle frontal gyrus (MFG: 3 contacts) and left orbitofrontal cortex (OF: 3 contacts) (Table 1). SPL-MFG and SPL-OF pairs showing NII_α_ of transitions from the two-stream to the one-stream percept (red line) and from the one-stream to the two-stream percept (blue line) in the endogenous **(B)** and exogenous **(C)** conditions. Purple dashed line marks the mean reaction time (1380 ms) of the exogenous (control) condition of the intracranial data. Statistical results are described in the main text.

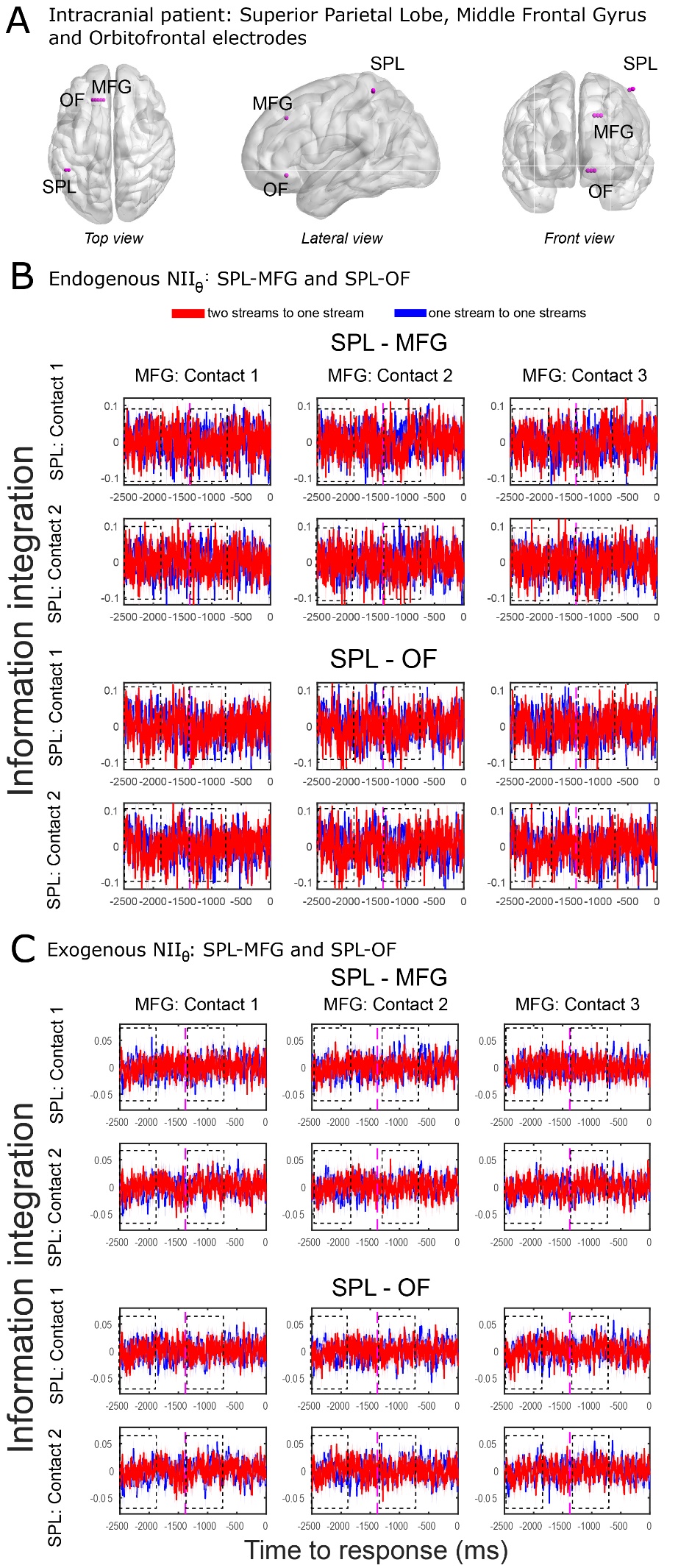

**Supplementary Figure 11. iEEG neural information integration in the theta range (NII_θ_).** **(A)** Electrodes were implanted in the left superior parietal lobe (SPL: 2 contacts), left middle frontal gyrus (MFG: 3 contacts) and left orbitofrontal cortex (OF: 3 contacts) (Table 1). SPL-MFG and SPL-OF pairs showing NII_θ_ of transitions from the two-stream to the one-stream percept (red line) and from the one-stream to the two-stream percept (blue line) in the endogenous **(B)** and exogenous **(C)** conditions. Purple dashed line marks the mean reaction time (1380 ms) of the exogenous (control) condition of the intracranial data. RANOVA revealed no statistical triple interaction between condition x window x direction (*F*_(1,56)_ = 0.58; *P* = 0.449).

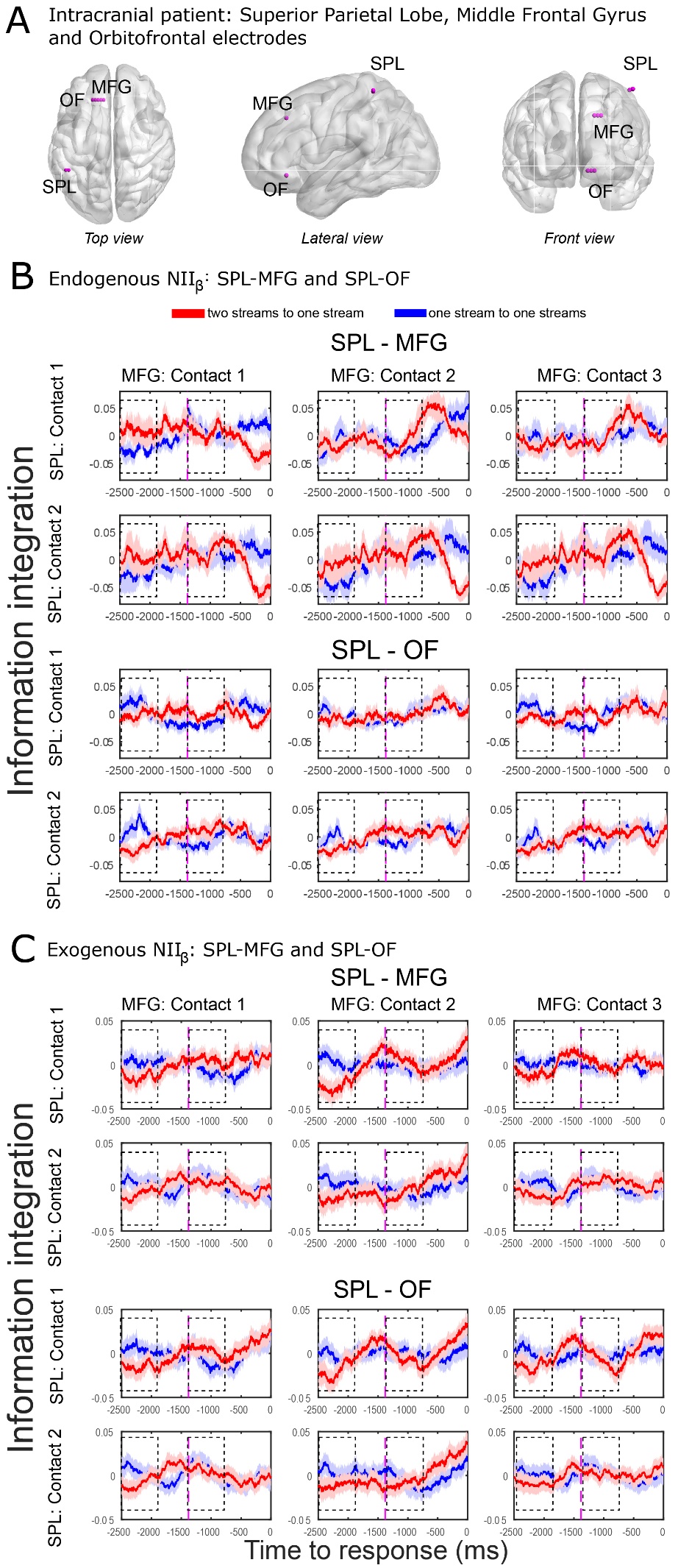

**Supplementary Figure 12. iEEG neural information integration in the beta range (NII_β_).** **(A)** Electrodes were implanted in the left superior parietal lobe (SPL: 2 contacts), left middle frontal gyrus (MFG: 3 contacts) and left orbitofrontal cortex (OF: 3 contacts) (Table 1). SPL-MFG and SPL-OF pairs showing NII**_β_** of transitions from the two-stream to the one-stream percept (red line) and from the one-stream to the two-stream percept (blue line) in the endogenous **(B)** and exogenous **(C)** conditions. Purple dashed line marks the mean reaction time (1380 ms) of the exogenous (control) condition of the intracranial data. RANOVA revealed no statistical triple interaction between condition x window x direction (*F*_(1,56)_ = 0.58; *P* = 0.475).

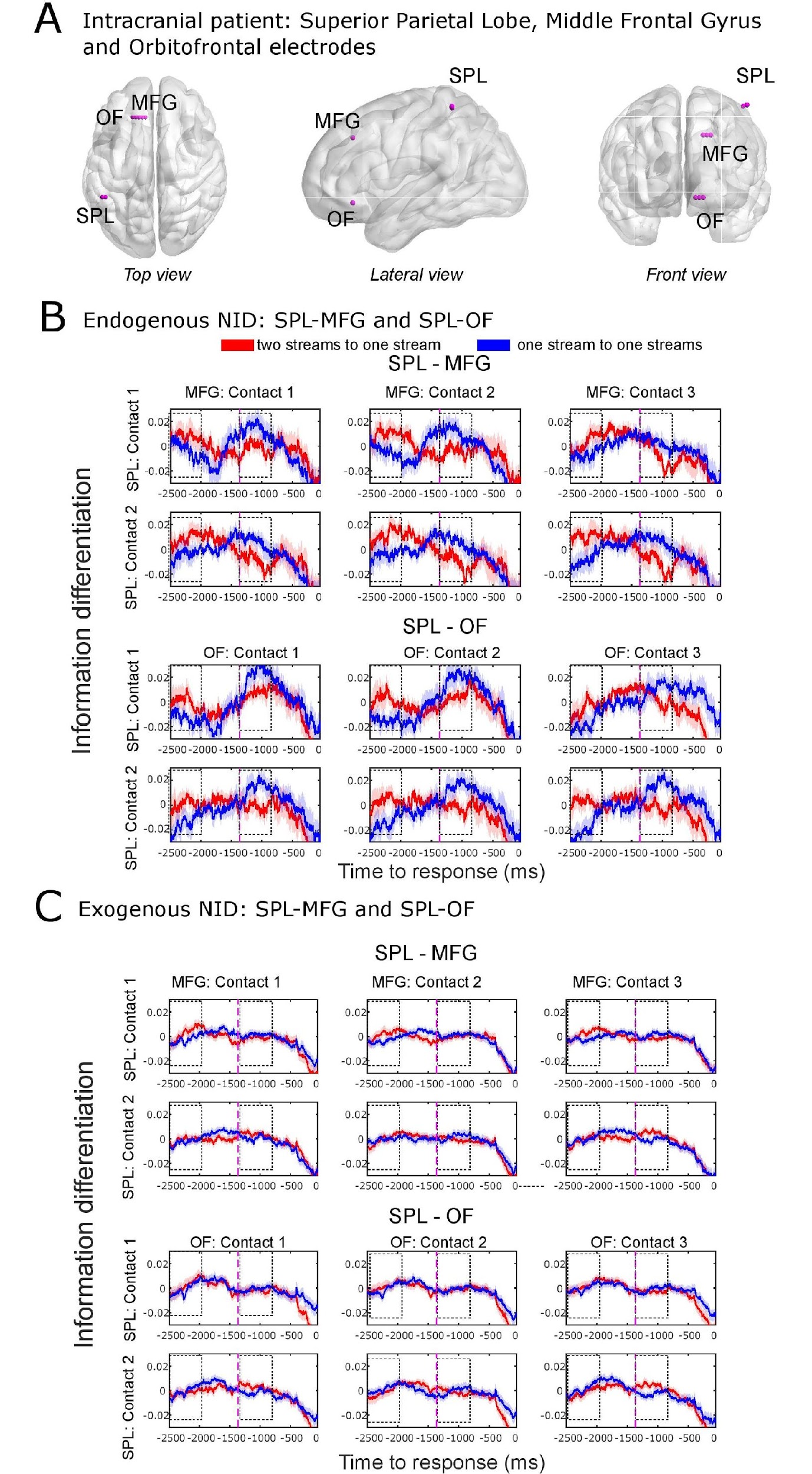

**Supplementary Figure 13. iEEG neural information differentiation (NID).** **(A)** Electrodes were implanted in the left superior parietal lobe (SPL: 2 contacts), left middle frontal gyrus (MFG: 3 contacts) and left orbitofrontal cortex (OF: 3 contacts) (Table 1). SPL-MFG and SPL-OF pairs showing NII_α_ of transitions from the two-stream to the one-stream percept (red line) and from the one-stream to the two-stream percept (blue line) in the endogenous **(B)** and exogenous **(C)** conditions. Purple dashed line marks the mean reaction time (1380 ms) of the exogenous (control) condition of the intracranial data.

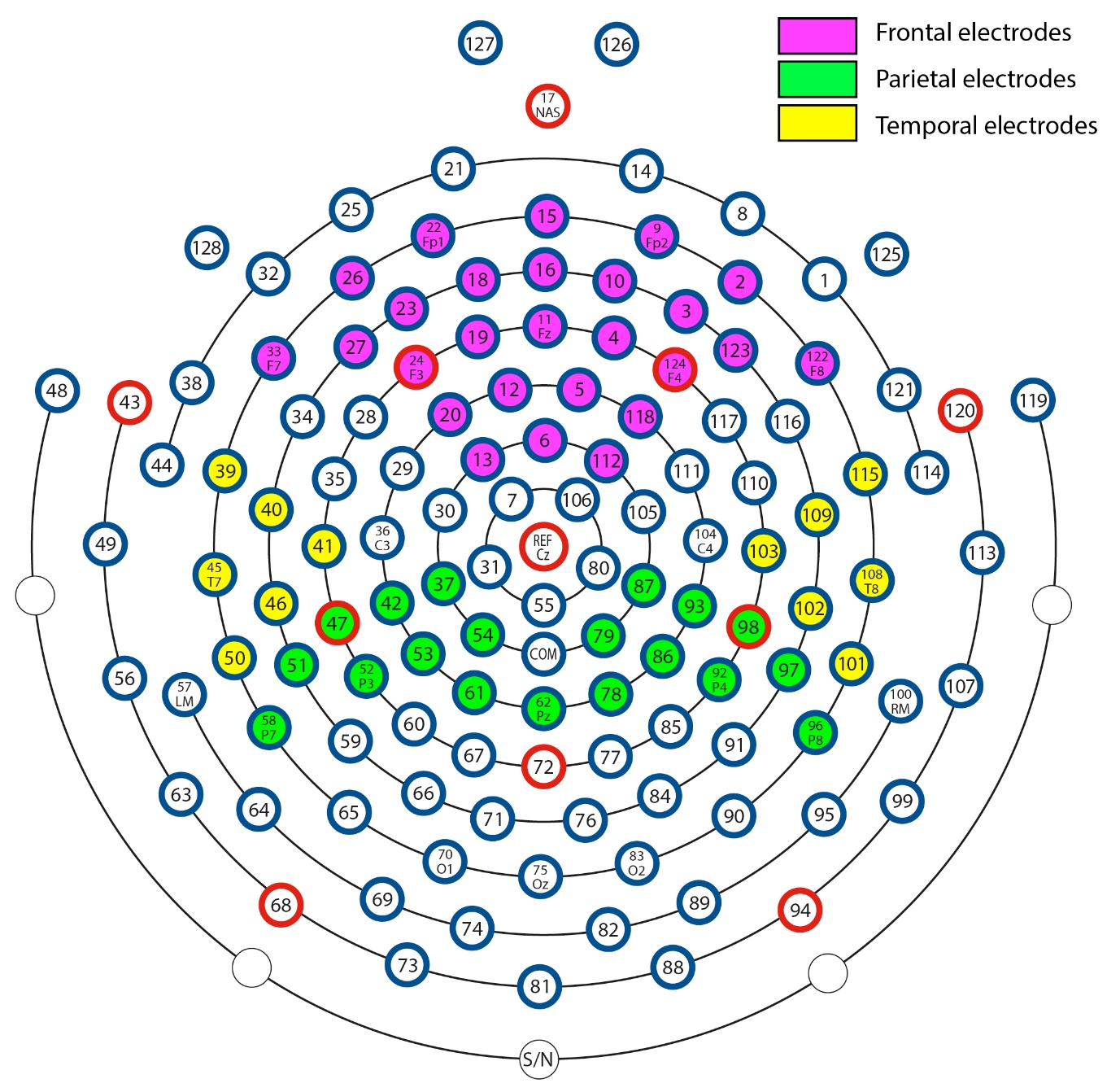

**Supplementary Figure 14.** Selected Frontal (green), Parietal (magenta) and Temporal (yellow) electrodes.
